# Preterm infants harbour diverse *Klebsiella* populations, including atypical species that encode and produce an array of antimicrobial resistance- and virulence-associated factors

**DOI:** 10.1101/761924

**Authors:** Yuhao Chen, Thomas C. Brook, Cho Zin Soe, Ian O’Neill, Cristina Alcon-Giner, Onnicha Leelastwattanagul, Sarah Phillips, Shabhonam Caim, Paul Clarke, Lindsay J. Hall, Lesley Hoyles

## Abstract

*Klebsiella* spp. are frequently enriched in the gut microbiota of preterm neonates, and overgrowth is associated with necrotizing enterocolitis, nosocomial infections and late-onset sepsis. Little is known about the genomic and phenotypic characteristics of preterm-associated *Klebsiella* as previous studies have focussed on recovery of antimicrobial-resistant isolates or culture-independent molecular analyses. Faecal samples from a UK cohort of healthy and sick preterm neonates (*n*=109) were screened on MacConkey agar to isolate lactose-positive *Enterobacteriaceae*. Whole-genome sequences were generated for isolates. Approximately one-tenth of faecal samples harboured *Klebsiella* spp. (*Klebsiella pneumoniae*, 7.3 %; *Klebsiella quasipneumoniae*, 0.9 %; *Klebsiella grimontii*, 2.8 %; *Klebsiella michiganensis*, 1.8 %). Isolates recovered from NEC- and sepsis-affected infants and those showing no signs of clinical infection (i.e. ‘healthy’) encoded multiple β-lactamases, which may prove problematic when defining treatment regimens for NEC or sepsis, and suggest ‘healthy’ preterm infants contribute to the resistome. No difference was observed between isolates recovered from ‘healthy’ and sick infants with respect to *in vitro* siderophore production (all encoded enterobactin in their genomes). All *K. pneumoniae*, *K. quasipneumoniae*, *K. grimontii* and *K. michiganensis* faecal isolates tested were able to reside and persist in macrophages, indicating their immune evasion abilities. Using a curated dataset of *Klebsiella oxytoca*, *K. grimontii* and *K. michiganensis* whole-genome sequences, metapangenome analyses of published metagenomic data confirmed our findings regarding the presence of *K. michiganensis* in the preterm gut, and highlight the importance of refined analyses with curated sequence databases when studying closely related species present in metagenomic data.

## INTRODUCTION

The gut microbiota encompasses bacteria, archaea, lower eukaryotes and viruses, with these microbes contributing to host gastrointestinal (GI) and systemic health. Host–microbiome interactions within the intestine are particularly important in neonates, contributing to development of the immune response, establishment of the gut microbiome and protection from infections [1, 2]. Term infants (i.e. gestation 37 weeks) are rapidly colonised after exposure to the mother’s microbiota and the environment, with streptococci and *Enterobacteriaceae* dominating in the initial phases [1], and *Bifidobacterium* spp. becoming prominent as the infant grows [1].

In contrast, colonization of preterm infants (i.e. <37 weeks’ gestation) occurs in neonatal intensive care units (NICUs) and is shaped by the significant number of antibiotics (‘covering’ (i.e. to cover possible early onset infection from birth) and treatment) these infants receive in the first days and weeks post birth. The microbiota in preterm infants is enriched for bacteria such as *Enterobacteriaceae*, *Enterococcus* and *Staphylococcus* [3, 4].

Critically, colonization of these at-risk infants with potentially pathogenic taxa, in concert with an unstable microbiome, and immaturity of their GI tract and immune system, is thought to contribute to nosocomial infections such as late-onset sepsis (LOS) or necrotizing enterocolitis (NEC) [5–10].

The family *Enterobacteriaceae* is a large group of Gram-negative bacteria encompassing many pathogens [e.g. *Escherichia* (*Esc.*) *coli*, *Klebsiella pneumoniae*, *Shigella* (*Shi.*) *dysenteriae*, *Enterobacter* (*Ent.*) *cloacae*, *Serratia* (*Ser.*) *marcescens* and *Citrobacter* spp.]. While coagulase-negative staphylococci are the most common cause of LOS in preterm infants, *Enterobacteriaceae* that translocate from the preterm gut to the bloodstream also cause this condition [8, 9]. In addition, *Enterobacteriaceae* are associated with higher morbidity than the staphylococci, and blooms in *Proteobacteria* – thought to be linked to impaired mucosal barrier integrity – have been reported immediately prior to the diagnosis of LOS [8,9,11]. Predictions made from shotgun metagenomic data show replication rates of all bacteria – and especially the *Enterobacteriaceae* and *Klebsiella* – are significantly increased immediately prior to NEC diagnosis [12]. This altered gut microbiome influences intestinal homeostasis and contributes to NEC [13], in tandem with the immature preterm immune system contributing to intestinal pathology in response to blooms of *Proteobacteria*.

Associations between *Klebsiella*-related operational taxonomic units (OTUs) and the development of NEC have been noted, suggesting members of this genus contribute to the aetiology of NEC in a subset of patients [14, 15]. Although Sim *et al.* [14] found one of their two distinct groups of NEC infants had an overabundance of a *Klebsiella* OTU, these researchers failed to identify a single predominant species of *Klebsiella*, recovering representatives of several genera (*K. pneumoniae*, *Klebsiella oxytoca*, *Ent. cloacae*, *Ent. aerogenes*, *Esc. coli* and *Ser. marcescens*) from samples. *Klebsiella* spp. and their fimbriae-encoding genes were significantly enriched in faeces collected immediately prior to the onset of NEC in a US infant cohort. These fimbriae may contribute to the overexpression of TLR4 receptors observed in preterm infants [12]. Confirming the role of these bacteria in NEC will require reproducing certain aspects of the disease in model systems, using well-characterized bacteria recovered from preterm infants [11, 14].

To date, there is limited information on the genomic and phenotypic features of preterm-associated *Klebsiella* spp. Thus, to characterise these important opportunistic pathogens, and to build a collection of preterm-associated *Klebsiella* strains for use in future mechanistic studies relevant to preterm-infant health, we isolated and characterized (phenotypically and genomically) bacteria from a cohort of preterm neonates enrolled in a study at the Norfolk and Norwich University Hospital (NNUH), Norwich, United Kingdom. Recovered *Klebsiella* isolates were subject to additional phenotypic tests that complemented genomic data. In addition, for the increasingly important species *K. oxytoca*, metapangenome analyses were undertaken to better understand the prevalence and potential virulence of this organism and related species in the context of the preterm neonate gut microbiota.

## METHODS

### Collection of faecal samples

Faeces were collected from premature neonates (<37 weeks’ gestation) (**Supplementary Table 1**). The Ethics Committee of the Faculty of Medical and Health Sciences of the University of East Anglia (Norwich, UK) approved this study. The protocol for faeces collection was laid out by the Norwich Research Park (NRP) Biorepository (Norwich, UK) and was in accordance with the terms of the Human Tissue Act 2004 (HTA), and approved with licence number 11208 by the Human Tissue Authority. Infants admitted to the NICU of the NNUH were recruited by doctors or nurses with informed and written consent obtained from parents. Collection of faecal samples was carried out by clinical researchers and/or research nurses, with samples stored at -80 °C prior to DNA extraction.

### 16S rRNA gene sequencing and analyses

DNA was extracted from samples using the FastDNA SPIN Kit for Soil (MP Biomedicals) and processed for sequencing and analyses as described previously [16]. This 16S rRNA gene sequence data associated with this project have been deposited at DDBJ/ENA/GenBank under BioProject accession PRJEB34372.

### Isolation of bacteria and biochemical characterization

For isolation work, a single faecal sample from each baby (*n*=109; **Supplementary Table 1**) was thawed and 0.1 g homogenised in 1 mL TBT buffer (100 mM Tris/HCl, pH 8.0; 100 mM NaCl; 10 mM MgCl_2_•6H_2_O). Homogenates were serially diluted 10^-1^ to 10^-4^ in TBT buffer. Aliquots (50 µL) of homogenate were spread on MacConkey agar no. 3 (Oxoid Ltd) plates in triplicate and incubated aerobically at 37 °C overnight.

Differential counts (based on colony morphology) of all lactose-positive (i.e. pink) colonies were made in triplicate to calculate colony-forming units (CFUs) per gramme wet-weight faeces. One of each colony type per plate was selected and re-streaked on MacConkey agar three times to purify, incubating aerobically at 37 °C overnight each time. A single colony from each pure culture was re-suspended in 5 mL of sterile distilled water; the API 20E kit (bioMérieux) was used according to the manufacturer’s instructions to give preliminary identities for each of the isolates recovered.

### DNA extraction, whole-genome sequencing and assembly

DNA was extracted using a phenol– chloroform method fully described previously [17] from overnight cultures of strains, and sequenced using the 96-plex Illumina HiSeq 2500 platform to generate 125 bp paired-end reads [18]. Raw data provided by the sequencing centre were checked using fastqc v0.11.4 (https://www.bioinformatics.babraham.ac.uk/projects/fastqc/); no adapter trimming was required, and reads had an average Phred score >25. MetaPhlAn2.6 [19] was used to identify which species genome sequences represented. According to the results given by MetaPhlAn2.6, appropriate reference genomes were retrieved from Ensembl Genome (http://bacteria.ensembl.org/index.html) to guide reference-based assembly using BugBuilder v1.0.3b1 (default settings for Illumina data) [20]. Summary statistics for the *Klebsiella* genome sequences generated in this study, including accession numbers, can be found in **Supplementary Table 2**. This Whole Genome Shotgun project has been deposited at DDBJ/ENA/GenBank under BioProject accession PRJNA471164.

#### Genome analyses

Average nucleotide identity (ANI) between genome sequences of isolates and reference strains (*Klebsiella grimontii* 06D021^T^, GCA_900200035; *K. oxytoca* 2880STDY5682490, GCA_900083895.1; *Klebsiella michiganensis* DSM 25444^T^, GCA_002925905) was determined using FastANI (default settings) [21].

*K. oxytoca*, *K. michiganensis* and *K. grimontii* genomes were uploaded to the *Klebsiella oxytoca* MLST website (https://pubmlst.org/koxytoca/) sited at the University of Oxford [22] on 28 July 2019 to determine allele number against previously defined house-keeping genes (*rpoB*, *gapA*, *mdh*, *pgi*, *phoE*, *infB* and *tonB*). *K. pneumoniae* genomes were analysed using the Institut Pasteur MLST database (https://bigsdb.pasteur.fr/klebsiella/klebsiella.html). Kleborate [23, 24] and Kaptive [25] were used to identify capsular type and O antigen type.

Virulence genes were identified by BLASTP of genome amino acid sequences against the Virulence Factors of Pathogenic Bacteria Database (VFDB; ‘core dataset’ downloaded 27 July 2019) [26]; results are reported for >70 % identity and 90 % query coverage. Antimicrobial resistance (AMR) genes were identified by BLASTP against the Comprehensive Antibiotic Resistance Database (CARD) download (27 July 2019; protein homolog dataset) [27]; only strict and perfect matches with respect to CARD database coverage and bit-score cut-off recommendations are reported.

Genomic traits were visualized using anvi’o-5.5 [28] according to the pangenomic workflow. Briefly, for each figure presented herein, genomes were used to create an anvi’o contigs database, which contained ORFs predicted using Prodigal v2.6.3 [29]. A multiple-sequence alignment was created using BLASTP. Markov CL algorithm [30] was used to identify gene clusters (--mcl-inflation 10; high sensitivity for identifying gene clusters of closely related species or strain level). Gene clusters and genomes were organized using Euclidean distance and Ward linkage, with results visualized using GoogleChrome.

### Phenotypic characterization of *Klebsiella* isolates

#### Iron assays

Pre-cultures of *Klebsiella* isolates (5 mL) were grown overnight in LB broth (37 °C, 160 rpm). Aliquots (500 µL) were harvested (4000 rpm, 20 min) and the cell pellets washed twice with PBS. The cell suspensions (50 µL) were used to inoculate 5 mL cultures containing M9 minimal medium (Na_2_HPO_4_, 6.9 g/L; KH_2_PO_4_, 3 g/L; NaCl, 0.5 g/L; NH_4_Cl, 1 g/L; CaCl_2_, 0.1 mM; MgSO_4_, 2 mM; 0.2 % glucose) at 37 °C. At 20 h, bacterial growth and siderophore production were measured using the CAS assay [31]. An aliquot (100 µL) of the cell culture supernatant was mixed with CAS dye (100 µL), followed by the shuttle solution (4 µL) and siderophore production monitored at 620 nm at 4 h using a BioRad Benchmark Plus microplate spectrophotometer. A decrease in the blue colour of the CAS dye was measured using uninoculated medium as control. The estimated amount of total siderophore produced by *Klebsiella* isolates was calculated using the CAS standard curve based upon a desferrioxamine B standard (1:1).

#### Macrophage assays

All strains were grown on LB broth + 1.5 % agar and incubated overnight at 37 °C. THP-1 monocytes were obtained from ATCC (TIB-202) and were maintained in RPMI (Gibco: 72400021) plus 10 % heat-inactivated foetal bovine serum (FBS; Gibco: 10500064) in a humidified incubator at 37 °C with 5 % CO_2_. THP-1 monocytes were differentiated into macrophages in RPMI + 10 mM HEPES + 10 % FBS + 10 ng/mL phorbol 12-myristate 13-acetate (PMA; Sigma) and seeded at 1×10^5^ cells per well of a 96-well tissue culture dish and incubated for 15 h. Overnight cultures of bacteria were diluted 1:100 into fresh LB broth and grown until mid-exponential phase. Bacteria were then washed twice with PBS and diluted to 1×10^7^ cfu/mL in RPMI + 10 mM HEPES + 10 % FBS and 100 µL of bacteria was added to each well. Plates were then centrifuged at 300 *g* for 5 min to synchronize infections. Bacteria–macrophage co-culture was incubated at 37 °C/5 % CO_2_ for 30 min to allow for phagocytosis. Cells were then washed three times in PBS and medium was replaced with above culture medium supplemented with 300 µg/mL gentamicin and 100 units/mL polymyxin B to eliminate extracellular bacteria. Cell were again incubated at 37 °C/5 % CO_2_ for 1.5 h. Cells were then washed three times with PBS and medium for cells for later time points was replaced with culture medium supplemented with 300 µg/mL gentamicin and incubated for a further 4.5 h. Intracellular bacterial load was enumerated by lysing macrophages in PBS + 1% Triton X-100 for 10 min at room temperature, serially diluting cultures and plating on LB agar. Plates were incubated overnight at 37 °C and colonies counted the following day.

#### Calculation of antibiotic minimal inhibitory concentration (MIC) for the *Klebsiella* isolates

Broth microdilution method was used to calculate the MIC of the *Klebsiella* isolates. Serial two-fold dilutions of benzylpenicillin, gentamicin, and meropenem were added to sterile nutrient broth. The antibiotics used in this assay were supplied by the NICU of NNUH. Inoculum for each of the isolates was prepared using 10 mL from a fresh overnight culture. Microplates were incubated for 24 h at 37 °C under aerobic conditions. Optical density was monitored using a plate reader (BMG Labtech, UK) at 595 nm. MICs were determined as the lowest concentration of antibiotic inhibiting any bacterial growth. All experiments were repeated in triplicate. For the aminoglycoside gentamicin and the carbapenem meropenem, *Klebsiella* (*Enterobacteriaceae*) breakpoints were determined according to European Committee on Antimicrobial Susceptibility Testing (EUCAST) guidelines (version 8.1, published 16 May 2018, http://www.eucast.org/fileadmin/src/media/PDFs/EUCAST_files/Breakpoint_tables/v_8.1_Breakpoint_Tables.pdf). No EUCAST data were available for benzylpenicillin (EUCAST states this aminopenicillin has no clinically useful activity against *Enterobacteriaceae*).

#### Estimation of abundance of *K. oxytoca* in shotgun metagenomic data

We chose to analyse a published preterm gut metagenome dataset [32] in this study, as it had been previously used to identify associations between uropathogenic *Esc. coli* and NEC. Trimmed, human-filtered, paired-end read data deposited in the Sequence Read Archive by Ward *et al.* [32] are available under BioProject accession number 63661. Information on Ward samples included in this study can be found in **Supplementary Table 3**. Ward *et al.* [32] used MetaPhlAn to determine abundance of bacteria in samples. However, the marker genes used to enumerate *K. oxytoca* in the MetaPhlAn2.6 database are derived from 11 genomes, five of which are not *K. oxytoca* (*Raoultella ornithinolytica* 10-5246, GCF_000247895; *K. michiganensis* E718, GCF_000276705; *K. michiganensis* Kleb_oxyt_10-5250_V1, GCF_000247915; *K. michiganensis* KCTC 1686, GCF_000240325; *K. michiganensis* Kleb_oxyt_10-5242_V1, GCF_000247835). Therefore, relative abundance of bacteria was instead determined using Centrifuge [33]. While MetaPhlAn2.6 relies on a pre-compiled database of unique marker genes for determining taxonomic abundance, the Centrifuge database can be updated at will using genomes downloaded from NCBI. A bacteria- and archaea-specific complete genome database was generated for use with Centrifuge via NCBI on 1 July 2018. Species-level abundances, based on read-level data, for *K. oxytoca* and *K. michiganensis* in the study of Ward *et al.* [32] were determined. (NB: *K. grimontii* genomes were not included in the Centrifuge database, nor are they included in MetaPhlAn2.6 or the most-recent version of Kraken2.)

#### Metapangenome analyses of *K. oxytoca, K. michiganensis* and *K. grimontii*

A total of 162 *K. oxytoca*-related whole-genome sequences were retrieved from GenBank on 31 May 2018 (**Supplementary Table 4**). On the basis of *bla*_OXY_, phylogenetic and ANI analyses [34], these had been confirmed to belong to *K. oxytoca* (*n*=64), *K. grimontii* (*n*=24) and *K. michiganensis* (*n*=74) (**Supplementary Table 5**). Prokka v1.13.3 [35] was used to annotate the 162 downloaded and five infant genomes. The resulting .gff files were subject to pangenome analyses using Roary v3.12.0 (default settings) [36]. Genes present in 165–167 strains were defined as the core cluster, while those present in 25–164 strains were defined as the accessory cluster. Remaining genes that only existed in single strains were classified into strain-specific clusters. FastTree v2.1.10 [37] was used to generate a phylogenetic tree from the core gene alignment, with the tree visualized using FigTree v1.4.4 (http://tree.bio.ed.ac.uk/software/figtree/).

PanPhlAn (panphlan_pangenome_generation.py v1.2.3.6; panphlan_profile.py v1.2.2.3; panphlan_map.py v1.2.2.5; [38]) was used to profile strains within metagenomes using the Roary-generated pangenome dataset. Gene-family clusters across all 167 available genomes and centroid sequence files outputted from Roary were uploaded to PanPhlAn to build a Bowtie2-indexed pangenome database, against which raw reads (concatenated read pair files) were mapped using Bowtie2 v2.3.0. The coverages of all gene positions were detected and extracted using samtools v1.4.1, and then integrated to a gene-family coverage profile for each sample. Pangenome-related strains were predicted to exist if a consistent and similar coverage depth across a set of 5774 gene families was detected under the non-default parameters of PanPhlAn (--min_coverage 1, --left_max 1.70, --right_min 0.30; panphlan_profile.py). Principal component analysis (PCA) was performed on 500 accessory genes randomly selected from the pangenome using the R package FactoMineR [39], allowing us to distinguish different species at the gene level.

#### Recovery of MAGs from metagenomes

For metagenome samples in which *K. oxytoca*-related strains were identified using PanPhlAn, attempts were made to recover them as MAGs. All reads in samples were mapped against the pangenome database using Bowtie2 and mapped paired-end reads were extracted by using FastQ Screen v0.11.3 as new fastq files (with parameter --tag, --filter 3). The extracted paired-end reads were assembled using SPAdes v3.12.0 [40]. These assemblies were known as original MAGs. A genome size of 5.5 Mb was set as a strict threshold: any assembly whose genome size was lower than this threshold was not considered in downstream analyses. FastANI was applied to calculate the ANI (cut-off 95 %) between MAGs and the three species reference genomes to double-check the predominant species in corresponding samples. The quality of each MAG was assessed using CheckM v1.0.18 [41].

A small number of the original MAGs were of high quality [42], but some contained a large number of contaminant contigs. All original MAGs were decontaminated as follows. Coding sequences of species-specific genomes in the pangenome were predicted using Prodigal v2.6.3 (Hyatt et al., 2010) with default settings and resulting multi-FASTA files containing protein sequences were concatenated to single files, which were used to build *K. oxytoca*-, *K. michiganensis*- and *K. grimontii*-specific databases in Diamond [43] format. Contigs of the original MAGs were aligned against the corresponding database under different minimum identity (%) cutoffs to report sequence alignments (with parameter --id 95, 96, 97, 98, 99 and 100). All unmapped contigs and contigs <500 nt in length were discarded from the original MAGs and the quality of new MAGs acquired was evaluated again using CheckM, to identify a Diamond BLAST identity threshold at which decontamination was effective while maintaining high genome completeness.

## RESULTS AND DISCUSSION

### Composition of the microbiota of preterm neonates

First faecal samples available after birth were collected from 109 hospitalized preterm infants (*n* = 50 female; *n* = 59 male) in the NICU of the NNUH (**Supplementary Table 1**; **Figure 1a**). On the basis of 16S rRNA gene sequencing data, *Enterobacteriaceae* were detected in the faeces of 42 (38.5 %) of the infants (**Figure 1b**).

**Figure 1.**
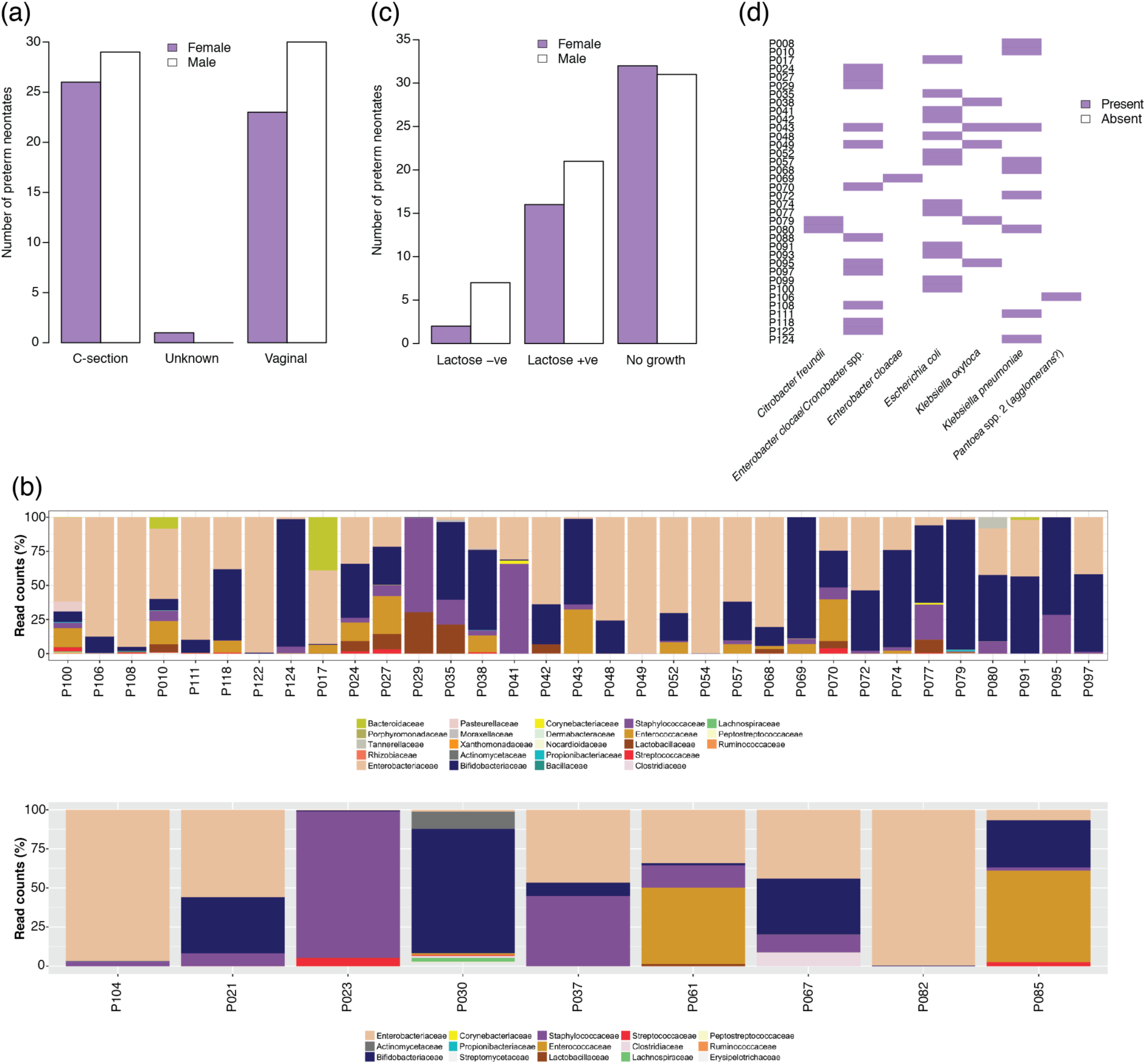
Summary information for UK cohort included in this study. (a) Breakdown of birth mode and sex of preterm neonates. (b) 16S rRNA gene sequence results for *Enterobacteriaceae*-positive samples: upper panel, samples from which lactose-positive isolates were recovered; lower panel, samples from which lactose-negative isolates were recovered. (c) Representation of lactose-negative and lactose-positive *Enterobacteriaceae* isolated from faecal samples. (d) Tentative identities of lactose-positive *Enterobacteriaceae* as determined by using API 20E.

All faecal samples were screened for *Enterobacteriaceae* using MacConkey agar no. 3 (**Figure 1c**). Forty-six (42.2 %) samples were positive for *Enterobacteriaceae* (*n* = 9 lactose-negative, carriage rate 8.3 %; *n* = 37 lactose-positive, carriage rate 33.9 %). Lactose-negative isolates were not characterized further. API 20E was used to provide tentative identification of LPE from 36 neonates (isolate could not be resuscitated for neonate P054) (**Figure 1d**). Of the 36 infants from whose faeces isolates were recovered, 23 were healthy, three had or were subsequently diagnosed with NEC, eight had suspected sepsis, one had an operation for gastroschisis and one was diagnosed with an eye infection after the faecal sample was taken (**Supplementary Table 1**).

### Whole-genome sequencing of neonatal faecal LPE

Whole-genome sequences were obtained for 56 LPE. MetaPhlAn2.6 was used to assign identities to genomes (not shown). Among the 56 isolates sequenced, 20 were identified as *K. pneumoniae* (carriage rate 8.3 %; **Supplementary Table 2**), 14 were *Ent. cloacae* complex (carriage rate 11.9 %), 13 were *Esc. coli* (carriage rate 11.9 %), five were *K. oxytoca* (**Supplementary Table 2**), two were *Citrobacter freundii* (carriage rate 1.8 %; [44]), one was *Citrobacter murliniae* (carriage rate 0.9 %; [44]) and one was *R. ornithinolytica* (carriage rate 0.9 %; [45]). *Esc. coli* and *Ent. cloacae* complex isolates will be discussed in detail elsewhere. MetaPhlAn2.6-generated identities matched those given by API 20E (**Supplementary Table 2**).

Reference-based assembly of genomes was performed using BugBuilder [20] (**Supplementary Table 2**). To determine whether preterm neonates may harbour more than one strain of a species in their faecal microbiota, nine isolates (#64–#73) were collected from neonate P008. These had all been identified as *K. pneumoniae* by API 20E and genome data. ANI across the nine isolates was >99.99 %. To determine whether the isolates were identical, gene content analysis was performed using Roary [36]. The average number of CDSs among these isolates was 5,385 (6.25) (**Supplementary Figure 1a–d**). Anvi’o showed the isolates were highly similar (**Supplementary Figure 1e**). Isolates of the same species from other neonates were also found to be identical to one another (#102 and #103 from P080; #118 and #119 from P124). For sets of identical isolates, only one was taken forward for further analyses. This left 14 distinct *Klebsiella* strains (9 *K. pneumoniae*; 5 *K. oxytoca*) for further analyses.

### Genome analyses of *K. pneumoniae* strains

*K. pneumoniae* is a commensal of the human gut microbiota and can cause nosocomial infections, NEC and LOS in premature neonates [9,14,46–49]. The genetic backgrounds of the neonate isolates were explored, to determine virulence and AMR genes encoded within the strains’ genomes.

Each isolate was genetically different: i.e. no two infants harboured the same strain of *K. pneumoniae* (**Supplementary Figure 2**). ANI analyses with representative strains of the seven phylogenetic groups of *K. pneumoniae* [50] showed eight of the neonatal isolates were *K. pneumoniae* (98.83–98.98 % ANI with *K. pneumoniae* ATCC 13883^T^ (GCA_000742135)) and one (#91) was *K. quasipneumoniae* (98.5 % ANI with *Klebsiella quasipneumoniae* subsp. *quasipneumoniae* 01A030^T^ (GCA_000751755)). MLST identified six STs within the *K. pneumoniae* strains (**Figure 2a**). *K. quasipneumoniae* #91 had a novel *mdh* allele, so no ST could be specified for this strain. None of the STs belonged to clonal complex (CC) 258, responsible for hospital outbreaks due to its frequent carriage of KPC and other acquired AMR genes [51]. The capsule of *K. pneumoniae* and related species is considered one of its major virulence factors. K1, K2 and K5 capsular types and hypervirulent types have strong associations with human infectious diseases [52, 53]. None of our neonatal isolates had a capsular type commonly associated with infections or hypervirulent *K. pneumoniae*, though K7, K10, K11, K16 and K38 isolates have previously been recovered from clinical samples in Taiwan [54]. Although the capsular type of strain #74 was identified as K62 with 99.33 % confidence and 100 % coverage, there was one gene (*KL62-12*, according to Kaptive) missing from the locus, leaving it a non-perfect match. Of the nine *K. pneumoniae* strains analysed using Kleborate [23, 24], O1v1 and O1v2 were represented equally among the O-antigen types (*n* =4 for both). These can be distinguished using genomic data but are serologically cross-reactive [24]. *K. quasipneumoniae* #91 was O3/O3a.

**Figure 2.**
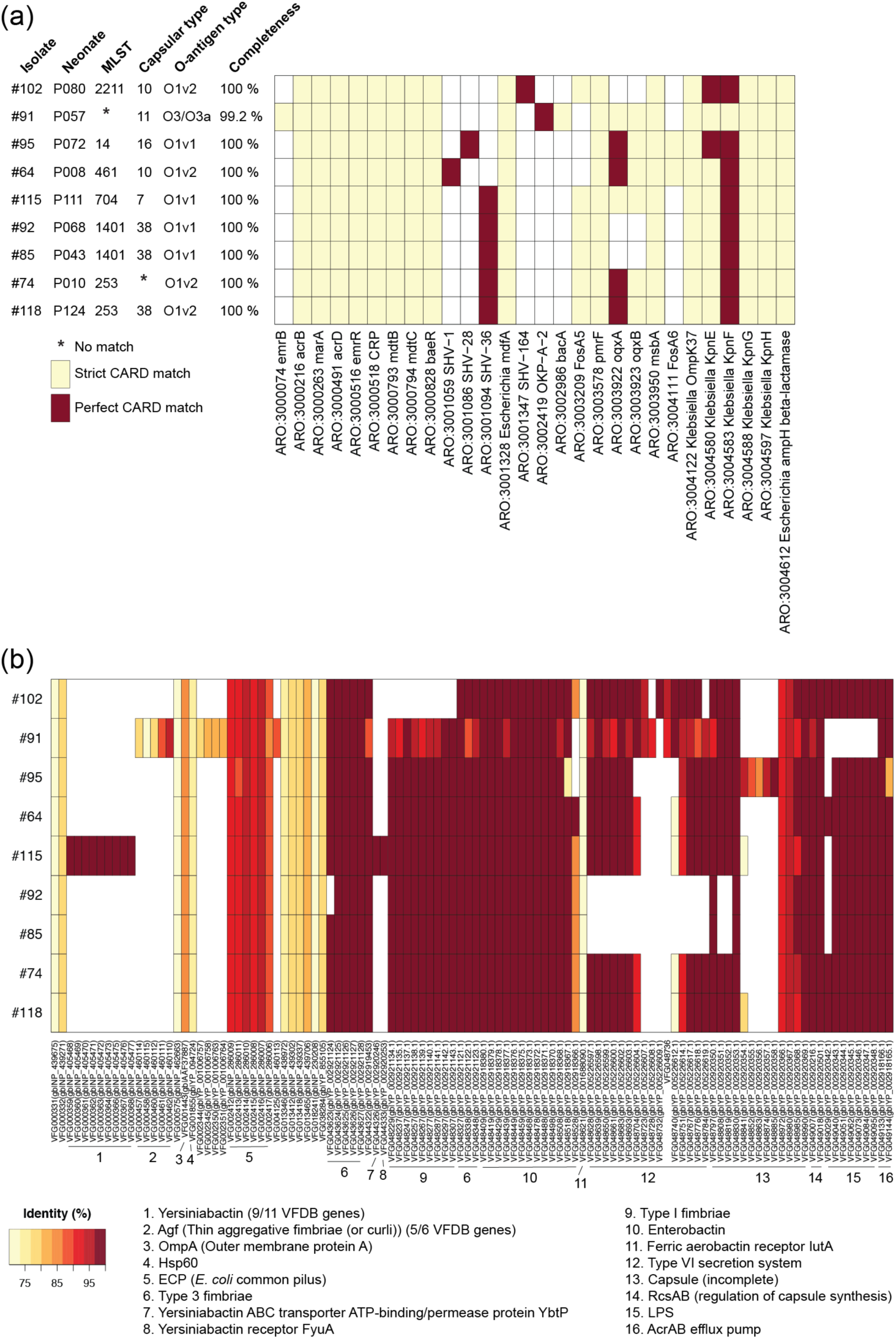
Summary of (a) antibiotic resistance and (b) virulence factor genes found in the *K. pneumoniae* isolates by comparison of protein sequences with those of the CARD and VFDB, respectively. (a) Strict CARD match, not identical but the bit-score of the matched sequence is greater than the curated BLASTP bit-score cut-off; perfect CARD match, 100 % identical to the reference sequence along its entire length. Loose matches are not shown to avoid presenting false positives based on sequences with low homology and bit-scores below CARD BLASTP cut-off recommendations. (b) Identity (%), BLASTP reported only for those proteins sharing >70 % identity and 90 % query coverage with VFDB protein sequences.

The vast majority (e.g. ∼90 % in the NNUH NICU) of preterm infants receive antibiotics during their NICU stay, often started routinely from admission (i.e. ‘covering’) if they are born very premature and/or very low birth weight. Administration of antibiotics can lead to disruption of early colonization by microbes, potentially encouraging growth of opportunistic pathogens such as LPE, creating a selection pressure that may promote development of AMR. All strains encoded homologues of *acrB*, *acrD*, *marA*, *emrR*, *CRP*, *mdtB*, *mdtC*, *baeR*, *Escherichia mdfA*, PmrF, *msbA*, OmpK37, KpnE, KpnF, KpnG, KpnH and *Escherichia ampH* β-lactamase, associated with antibiotic efflux and its regulation (*acrB*, *acrD*, *marA*, *emrR*, *CRP*, *mdtB*, *mdtC*, *baeR*) and resistance to: aminoglycosides; cationic antimicrobial peptides and antibiotics such as polymyxin (PmrF); chloramphenicol (*mdfA*), cefotaxime and cefoxitin (OmpK37); cefepime, ceftriaxone, colistin, erythromycin, rifampin, tetracycline, streptomycin as well as enhanced sensitivity toward sodium dodecyl sulfate, deoxycholate, dyes, benzalkonium chloride, chlorohexidine, and triclosan (KpnE, KpnF); azithromycin, ceftazidime, ciprofloxacin, ertapenem, erythromycin, gentamicin, imipenem, ticarcillin, norfloxacin, polymyxin-B, piperacillin, spectinomycin, tobramycin, and streptomycin (KpnG, KpnH); β-lactams and penicillin (*Escherichia ampH* β-lactamase). Homologues of *bla*_SHV_ were found in all *K. pneumoniae* strains except #91; a homologue of *bla*_OKP_ was encoded by *K. quasipneumoniae* #91, which also encoded homologues of *emrB* (a translocase that recognizes substrates including carbonyl cyanide *m*-chlorophenylhydrazone, nalidixic acid, and thioloactomycin and *bacA* (confers resistance to bacitracin) (**Figure 2a**). Plasmid-encoded *bla*_SHV_ enzymes represent an important subgroup of class A β-lactamases, while chromosomally encoded β-lactamase *bla*_OKP_ cannot hydrolyse extended-spectrum cephalosporins [55]. Homologues of *oqxA* and *oqxB* (encoding OqxAB, a plasmid-encoded efflux pump that confers resistance to fluoroquinolones) were encoded by #64, #74, #91, #95, #115 and #118. Strains #64 and #95 encoded homologues of FosA6 (confers resistance to fosfomycin), while #74, #85, #92, #115 and #118 encoded homologues of FosA5 (confers resistance to sulphonamide, cephalosporins, gentamicin, ciprofloxacin, chloramphenicol and streptomycin).

While the majority of the neonatal *K. pneumoniae* strains did not represent known pathogenic lineages, virulence factors were detected in their genomes (**Figure 2b**). The host limits iron availability within the GI tract to prevent colonization by pathogens and bacterial overgrowth. However, *Klebsiella* spp. have evolved numerous mechanisms to circumvent these defences. Thus, we determined whether gene clusters associated with iron uptake and siderophore systems (i.e. enterobactin, yersiniabactin, aerobactin, colibactin, salmochelin) were present in the strains. All strains encoded enterobactin, while only #115 encoded an additional system (yersiniabactin) (**Figure 2b**). All strains encoded *Esc. coli* common pilus, OmpA, Hsp60, type 3 fimbriae, ferric aerobactin receptor *IutA* and the AcrAB efflux pump. All strains except #102 encoded type 1 fimbriae; all strains except #91 encoded lipopolysaccharide (LPS). *K. quasipneumoniae* #91 encoded thin aggregative fimbriae, associated with biofilm formation and adhering to human mucosal or epithelial surfaces. Incomplete coverage of capsule genes is likely due to the limited database of VFDB compared with those used to populate Kaptive and Kleborate.

### Whole-genome analyses of isolates tentatively identified as *K. oxytoca*

*K. oxytoca* is a minor member of the human gut microbiota, recovered at low levels from the faeces of 1.6–9 % of healthy adults [56, 57]. Toxigenic *K. oxytoca* is a causative agent of antibiotic-associated haemorrhagic colitis, a condition affecting mainly young and otherwise healthy outpatients after brief treatment with penicillin derivatives [57]. *K. oxytoca* has been detected in the faeces of a subset of preterm infants via cultivation or shotgun metagenomics, but its association with preterm-associated infections is unknown [14,32,58]. At the DNA level, bacteria characterized phenotypically as *K. oxytoca* actually represent three phylogroups/distinct species: Ko1, *K. michiganensis*; Ko2, *K. oxytoca*; Ko6, *K. grimontii* [59–62]. *K. michiganensis* and *K. oxytoca* are distinguishable based on the *bla*_OXY_ gene they carry (*bla*_OXY-1_ and *bla*_OXY-2_, respectively) [59]. *K. grimontii* was recently described to accommodate Ko6 strains based on *rpoB*, *gyrA* and *rrs* gene sequences [61]. The colonization of humans with *K. oxytoca* phylogroups has previously been associated with the genetic backgrounds of strains: Ko2 mainly inhabits the lower GI tract, with Ko1 and Ko6 generally associated with respiratory isolates and faecal isolates, respectively [60].

On the basis of API 20E data and initial genome (MetaPhlAn2.6) analysis, five neonatal isolates were identified as *K. oxytoca* (#80, #83, #88, #99, #108). MetaPhlAn2.6 cannot distinguish among the three phylogroups. ANI of the genomes against reference genomes showed #88 and #108 to be *K. michiganensis* (both 98.78 and 98.94 % ANI, respectively with GCA_002925905) and #80, #83 and #99 to be *K. grimontii* (99.18, 99.23 and 99.17 % ANI, respectively, with GCA_900200035) **(Supplementary Figure 3a**), with ANI cut-off values well above the ∼95 % proposed for species delineation [63–65] and used by Passet & Brisse [61] to separate *K. grimontii* from *K. oxytoca* and *K. michiganensis*. Phylogenetic analysis with a panel of authentic *K. oxytoca*, *K. grimontii* and *K. michiganensis* genomes confirmed the species affiliations of the infant isolates (**Supplementary Figure 3b**). Similar to the *K. pneumoniae* isolates, no two infants harboured the same strain of *K. michiganensis* or *K. grimontii* (**Figure 3a**).

**Figure 3.**
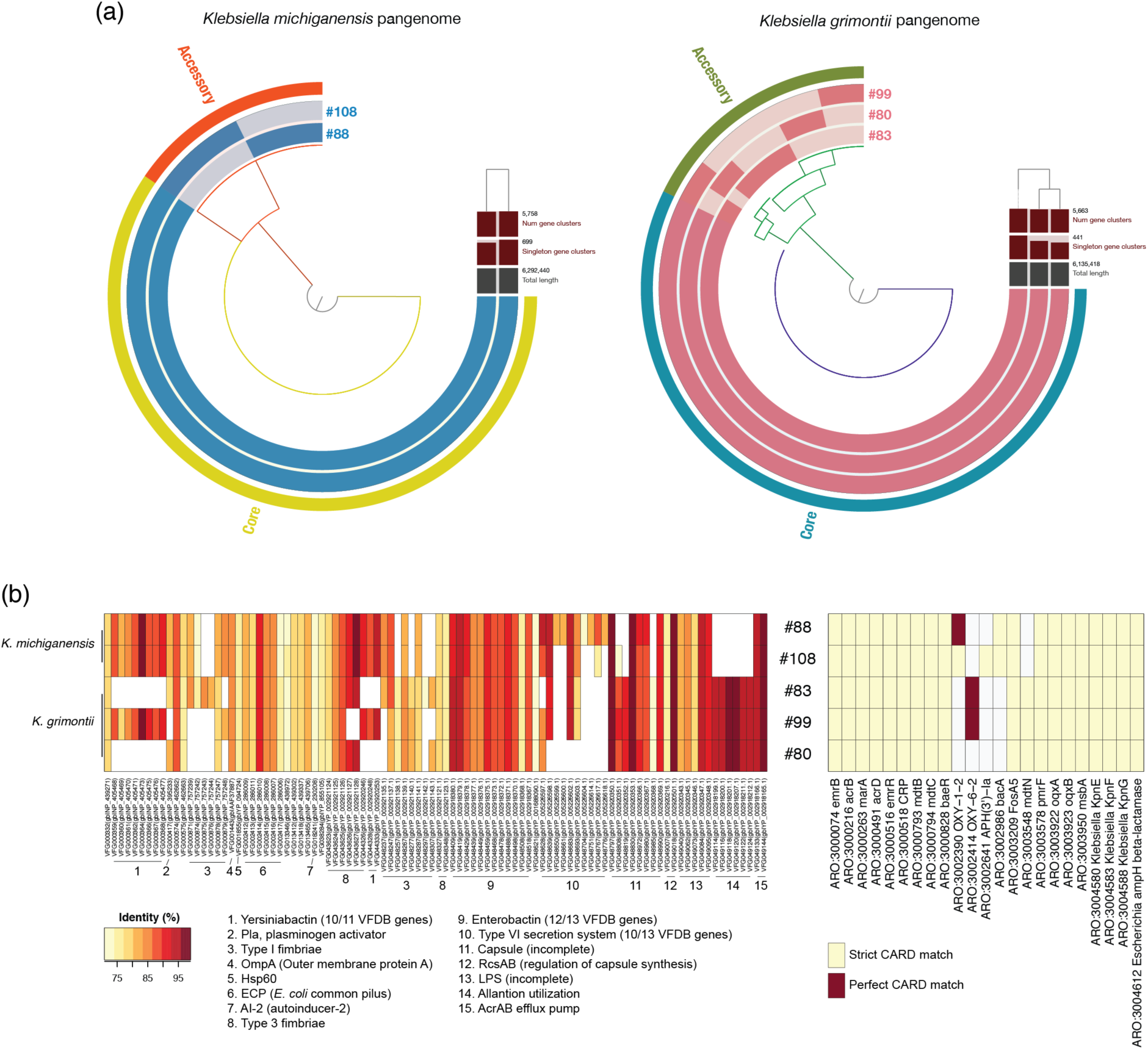
Genomic characterization of the *K. michiganensis* and *K. grimontii* isolates recovered from neonates. (a) Anvi’o representation of the genomes of *K. michiganensis* and *K. grimontii* isolates recovered from different infants. It is clear the isolates are different from one another at the genomic level. (b) Virulence factor (left side) and antibiotic resistance (right side) genes encoded by the isolates. Criteria for identity and strict/perfect match with respect to VFDB and CARD, respectively, are the same as those given for Figure 2.

### Predicted virulence and AMR determinant genes of infant-associated *K. michiganensis* and *K. grimontii*

The *K. michiganensis* and *K. grimontii* strains were examined for the presence of virulence-associated loci found in *K. pneumoniae* strains [51] (**Figure 3b**). Enterobactin was encoded by all strains. Yersiniabactin was predicted to be encoded by *K. grimontii* #99 and *K. michiganensis* #88 and #108. Other siderophore-associated gene clusters (aerobactin, colibactin and salmochelin) found in *K. pneumoniae* were absent. An allantoinase gene cluster (including *allB*/*C*/*R*/*A*/*S* and *ybbW*), which plays a role in *K. pneumoniae* liver infection [66], was identified in the three *K. grimontii* strains.

Due to the clinical importance of AMR in *Enterobacteriaceae*, an *in silico* AMR gene profile was established for the *K. michiganensis* and *K. grimontii* strains. Homologues of 18 AMR determinant genes (*emrB*, *emrR*, *acrB*, *acrD*, CRP, *marA*, *mdtB*, *mdtC*, *baeR*, FosA5, *pmrF oqxA*, *oqxB*, *msbA*, KpnE, KpnF, KpnG, *Escherichia ampH* β-lactamase) were common to the five strains, similar to the *K. pneumoniae* isolates. Both *K. michiganensis* strains encoded homologues of OXY-1-2 (β-lactamase specific to *K. michiganensis* (Ko1; [67]) and *bacA*, while #108 encoded a homologue of *aph(3’)-la* (aminoglycoside phosphotransferase). All *K. grimontii* strains encoded *mdtN* (potentially involved in resistance to puromycin, acriflavine and tetraphenylarsonium chloride), while #83 and #99 encoded homologues of OXY-6-2 (β-lactamase specific to *K. grimontii* (Ko6; [67]).

### Phenotypic characterization of *K. pneumoniae, K. quasipneumoniae, K. michiganensis* and *K. grimontii* neonatal isolates

Five of the 13 *Klebsiella* strains we characterised were isolated from preterm infants who had been diagnosed with either NEC or sepsis. Thus, we sought to link our genotypic analyses with clinically important virulence traits including the ability to survive and replicate in host immune cells (i.e. macrophages) and the ability to produce iron-acquiring siderophores. We also determined the strains’ AMR profiles.

Previous studies have indicated that respiratory infection-associated *K. pneumoniae* are able to survive within macrophages, a critical innate immune cell type required for optimal pathogen clearance [68]. However, to date there is limited information relating to this ability in gut-associated strains, and there is no information on other *Klebsiella* species. Thus, all *Klebsiella* strains isolated in this study were tested in PMA-differentiated THP-1 macrophages using a gentamicin protection assay. All strains appeared to persist within macrophages, as bacterial load was either maintained over the time-course or increased or decreased between 1.5 h and 6 h, although these values were not statistically significant (**Figure 4a**). These data suggest that all *Klebsiella* strains tested can reside and persist in macrophages. This ability of all strains to survive, and in some cases potentially replicate, within macrophages indicates their immune evasion capabilities, which may link to increased risk and incidence of NEC and sepsis if these strains translocate from the ‘leaky’ preterm GI tract to systemic sites contributing to the inflammatory cascades characteristic of these conditions.

**Figure 4.**
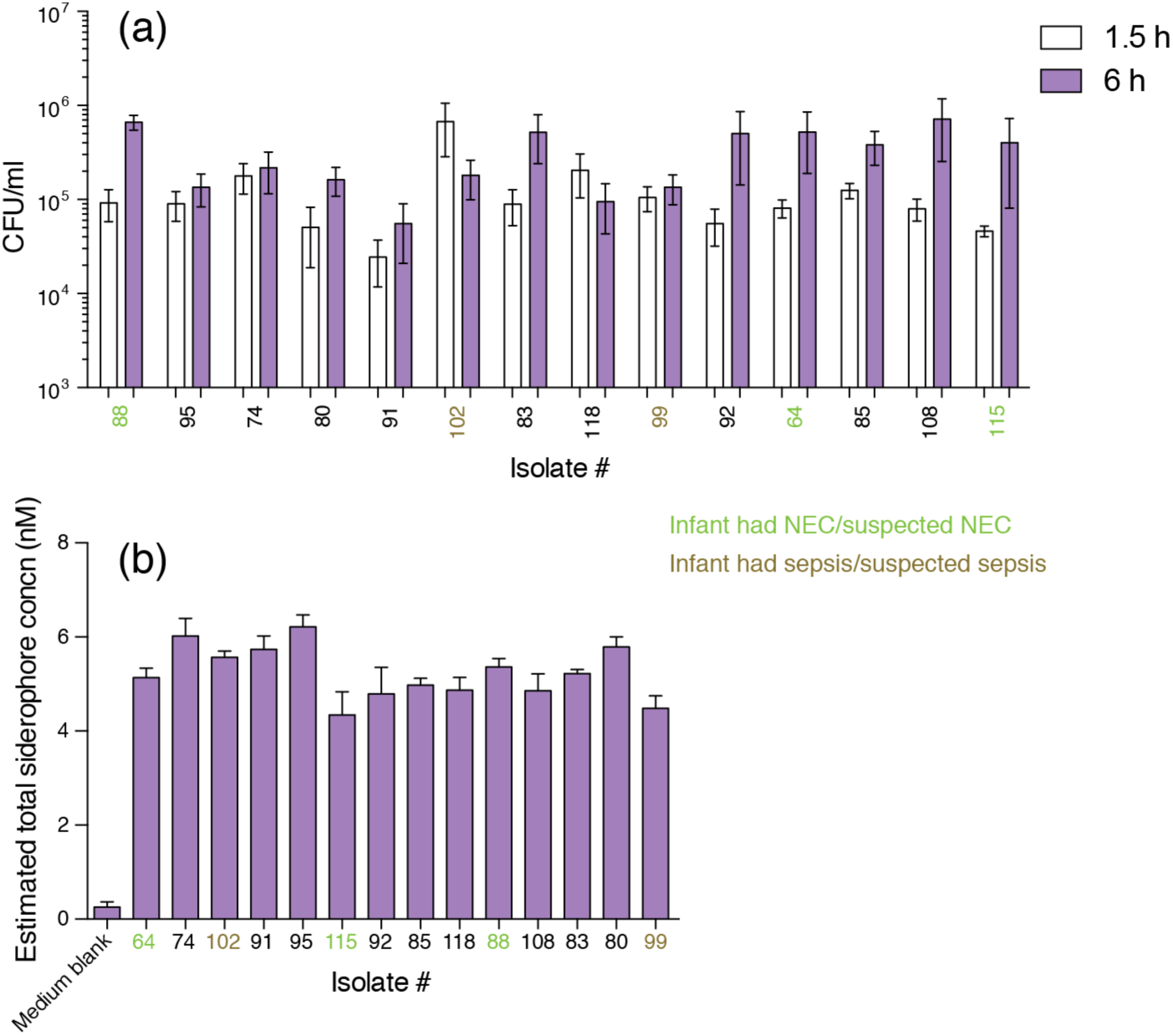
Phenotypic assays for the *Klebsiella* isolates recovered from infants. (a) Strains were tested for persistence in PMA-differentiated THP-1 macrophages using a gentamicin protection assay. Intracellular bacteria were enumerated 1.5 h and 6 h after infection to determine persistence (*n*=4). Results are shown as mean (SD). (b) *Klebseilla* strains were grown in minimal medium and at 20 h bacterial growth (OD_600_) siderophore production was measured using the CAS assay (*n* = 3). Results are shown as mean (SD).

Iron is a vital nutrient that performs multiple roles in cellular processes, ranging from DNA replication and cell growth to protection against oxidative stress. In the healthy host, the majority of iron is bound with intracellular proteins and the remaining free iron is extracellular and insoluble, hence difficult to access [69]. For invading pathogens, siderophore systems are critical for iron competition and uptake to accomplish colonization and cause infections, this is particularly true in the preterm GI tract. Preterm infants in NICU are heavily supplemented with iron as they receive many red-blood-cell transfusions (increasing hepatic iron stores), iron-supplemented parenteral nutrition, and supplementary oral iron within a few weeks of birth. During infection, *Klebsiella* secretes siderophores to sequester iron and to establish colonization in the host. Enterobactin is the most well-known siderophore produced by *K. pneumoniae* and related species, and was found to be encoded by all our strains (**Figure 2b**, **Figure 3b**). The host innate immune protein lipocalin 2 binds to enterobactin and disrupts bacterial iron uptake [70]. *Klebsiella* species have evolved to hoodwink this host response by producing several evasive siderophores [71, 72]. Siderophore production of *Klebsiella* isolates was monitored using CAS liquid assay. All isolates tested grown in M9 minimal medium were CAS-positive with the estimated siderophore concentration ranging between of 3.5 and 6 nM (**Figure 4b**). There was no significant difference in siderophore production between ‘healthy’ and NEC- and sepsis-associated isolates.

*Klebsiella* is of concern within an AMR context, particularly in at-risk neonates, due to the increasing emergence of multidrug-resistant isolates that cause severe infection [73]. A UK study in which 24 % of all LOS cases were caused by *Enterobacteriaceae* (8.9 % of all caused by *Klebsiella* spp.) showed a high proportion (14 % and 34 %, respectively) of *Enterobacteriaceae* isolates recovered from sick infants were resistant to flucloxacillin/gentamicin and amoxicillin/cefotaxime, the two most commonly used empiric antibiotic combinations [49]. Thus, to demonstrate antibiotic-resistance phenotypes in *Klebsiella* spp. correlating to presence of AMR genotypes, we tested the susceptibility of the isolates with three antibiotics commonly prescribed in NICUs; gentamicin, meropenem and benzylpenicillin (**Table 1**). One strain of *K. grimontii* (#80) was potentially sensitive to benzylpenicillin, an aminopenicillin currently not recognized as being clinically useful against *Enterobacteriaceae*. Isolates #64 (encoded KpnG and KpnH), #83 and #108 (both encoding FosA5) were resistant to gentamicin, while #80, #88 and #99 (all encoding FosA5) and #95 (encoding KpnG and KpnH) showed intermediate susceptibility to this aminoglycoside.

Presence of a gene in a bacterium’s genome does not mean it is functionally active, nor does it give any indication as to how active the gene is if it is indeed functional: e.g. all nine *K. pneumoniae* isolates encoded KpnG and KpnH, but only two showed any resistance to gentamicin upon susceptibility testing.

Isolates #64 (SHV-1), #74 (SHV-36), #85, #88, #91 (OKP-A-2), #92 (SHV-36), #95 (SHV-28), #102 (SHV-164), #115 (SHV-36) and #118 (SHV-36) – which all encoded β-lactamases (shown in parentheses in preceding text, along with *Escherichia ampH* β-lactamase) – showed intermediate susceptibility to the carbapenem meropenem. *K. pneumoniae* #64 was isolated from an infant with clinically diagnosed NEC with confirmed *Klebsiella* colonization. Importantly this preterm infant had previously been treated with benzylpenicillin, gentamicin and meropenem, which may link to the observed phenotypic resistance and corresponding AMR genes *Escherichia ampH* β-lactamase, KpnG/KpnH and SHV-1, respectively, and suggests further treatment with gentamicin and meropenem would have been ineffective in this infant. Indeed, the infant was treated with cefotaxime, metronidazole and vancomycin in a subsequent round of medication (**Supplementary Table 2**). *K. pneumoniae* #115 was isolated from a baby that had confirmed NEC: the strain was resistant to benzylpenicillin (encoded *Escherichia ampH* β-lactamase) and showed intermediate resistance to meropenem (encoded SHV-36). *K. michiganensis* #88, also isolated from a baby that had NEC, showed intermediate resistance to both benzylpenicillin (encoded FosA5) and meropenem (encoded OXY-1-2): both antibiotics had been administered to the baby at birth. *K. grimontii* #99, isolated from a baby with suspected sepsis, showed intermediate resistance to benzylpenicillin (encoded FosA5).These data indicate that preterm-associated *Klebsiella* have a multi-drug resistant phenotype that may prove problematic when treatment options are required for sepsis or NEC. Interestingly, other isolates (e.g. #95, recovered from an infant who had received benzylpenicillin and gentamicin; **Supplementary Table 2**) associated with ‘healthy’ preterm infants also harboured AMR genes (#95: KpnG, KpnH, SHV-28, *Escherichia ampH* β-lactamase) and phenotypic resistance profiles suggesting that administration of antibiotics to preterm infants with no signs of clinical infection contributes to the reservoir of AMR genes – the ‘resistome’ [74] – which may increase horizontal gene transfer of AMR determinants to other opportunistic pathogens residing within the GI tract.

### Abundance of *K. oxytoca* and related species in metagenomic datasets

We used a published metagenomics dataset [32] to determine the prevalence of *K. oxytoca*, *K. michiganensis* and *K. grimontii* in the preterm infant gut microbiome. These data had previously been used to look at the relationship between NEC and uropathogenic *Esc. coli*, and metadata were available for the samples. Ward *et al.* [32] collected a total of 327 samples at three stages of infant life: stage1, days 3–9; stage2, days 10–16; stage3, days 17–22. Within each life stage samples were collected on more than one day for some infants. In the current study, only samples processed under Protocol A of Ward *et al.* [32] and from the earliest collection day within each life stage were analysed. For those samples for which multiple sets of paired-end data were available, read data were concatenated and used in analyses (**Supplementary Table 3**).

Stage1 comprised samples from 127 infants (105 preterm, 22 term), 16 of whom had been diagnosed with NEC and 10 infants had subsequently died. Stage2 comprised samples from 146 infants (128 preterm and 18 term), 24 of whom later developed NEC with 18 deaths. Stage3 comprised samples from 54 infants (48 preterm, 6 term), including eight NEC patients, six of whom died. Samples were collected from 165 distinct infants (143 preterm, 22 term) but only 41 of them were sequenced at all three life stages. Infants were born either vaginally (*n* = 70) or by caesarean section (*n* = 95). The gestational ages of preterm infants ranged from 23 to 29 weeks (mean 26.1 weeks), while the term babies ranged from 38 to 41 weeks (mean 39.2 weeks). As we had found that the MetaPhlAn2.6 database contained non-*K. oxytoca* genomes within its *K. oxytoca* dataset (detecting *K. oxytoca*, *K. michiganensis* and *R. ornithinolytica* (refer to Methods)), we used Centrifuge to determine abundance of this species in metagenomes (**Figure 5ab**). Due to their genomic similarity, *K. oxytoca* and *K. michiganensis* could not be readily distinguished using Centrifuge (**Figure 5a**); no genomes assigned to *K. grimontii* were included in the Centrifuge database at the time this study was undertaken. Though it should be noted that, while Centrifuge (and Kraken2) relies on NCBI taxonomy for species identification, there are still many genomes within GenBank/RefSeq that are assigned to the wrong species (e.g. assemblies GCA_001052235.1 and GCA_000427015.1 within our curated pangenome dataset have been confirmed by detailed analyses to be *K. grimontii* (**Supplementary Figure 2** and [34]), but still assigned as *K. oxytoca* and *K. michiganensis*, respectively, within GenBank as of 28 July 2019; these are by no means the only examples from our current study).

**Figure 5.**
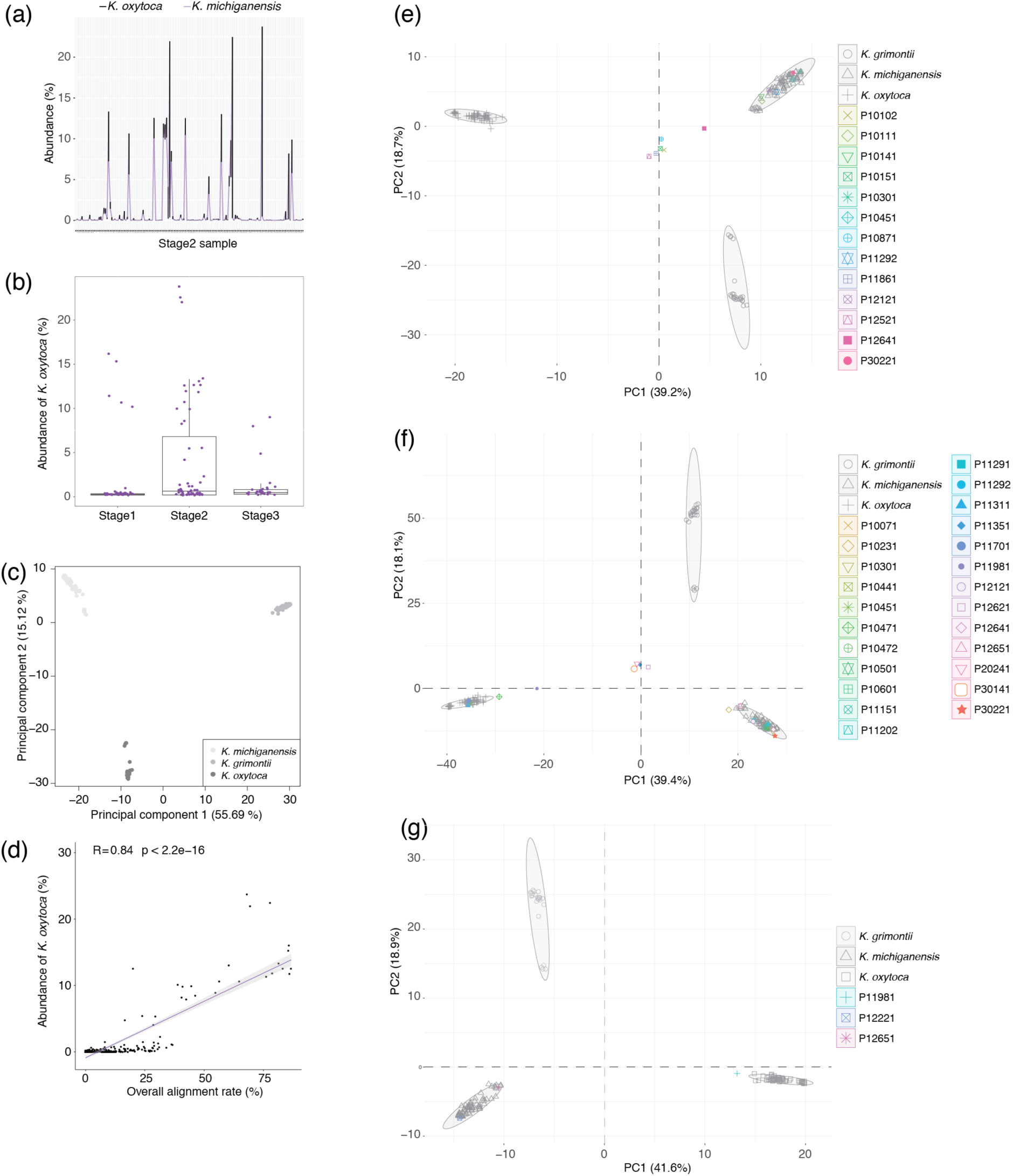
Identification of *K. oxytoca*-related species in infant faecal metagenomes. (a) Comparison of *K. oxytoca* and *K. michiganensis* abundance (as determined using Centrifuge) in stage2 samples of Ward *et al.* [32]. (b) Abundance of *K. oxytoca* (determined using Centrifuge) across stage1, stage2 and stage3 samples of Ward *et al.* [32]. (c) Separation of the strains of *K. grimontii* (*n*=27), *K. michiganensis* (*n*=76) and *K. oxytoca* (*n*=64) based on accessory genes (*n* = 5,108) detected in the Roary-generated open pangenome. (d) Relationship between PanPhlAn (overall alignment rate) and Centrifuge (abundance *K. oxytoca* (%)) data. (e, f, g) PCA plots show separation of strains in the pangenome plus PanPhlAn-detected strains based on presence of 500 randomly sampled accessory genes at (e) stage1, (f) stage2 and (g) stage3 of Ward *et al.* [32].

For those samples harbouring *K. oxytoca*, relative abundance of the bacterium increased from stage1 to stage2 and decreased at stage3 (**Figure 5b**).

### Metapangenome analysis of preterm infant metagenomic data to detect *K. oxytoca*, *K. michiganensis* and *K. grimontii*

Using a set of 162 *K. oxytoca*-related genomes (**Supplementary Table 4**) and those of the five infant isolates, a pangenome was generated using Roary. The pangenome dataset consisted of 76 *K. michiganensis* (mean ANI among strains 98.55 (0.60) %, range 97.13–100 %), 64 *K. oxytoca* (mean ANI among strains 99.20 (0.30) %, range 98.53–100 %) and 27 *K. grimontii* (mean ANI among strains 98.45 (1.47) %, range 95.70–100 %) strains. A total of 40,605 genes were detected in the open pangenome: 2,769 of them constituted the core gene cluster, while the accessory cluster included 5,108 genes and the remaining 32,728 genes formed the strain-specific cluster. A PCA plot based on the accessory genes clustered strains into the three different species (**Figure 5c**), in agreement with our phylogenetic analysis of the core genes (**Supplementary Figure 3b**) and consistent with the findings of [75], who were able to split the three species (phylogroups) based on a pangenome analysis of fewer genomes.

The Roary-generated pangenome was used as a custom database for PanPhlAn, to detect the presence and absence of core genes and accessory genes in each infant sample. As expected, the proportion of reads that PanPhlAn mapped to the custom database correlated with the Centrifuge-generated abundance data (**Figure 5d**). In stage1, 13 infants (12 preterm, 1 term) were predicted to carry *K. oxytoca*-related species (**Figure 5e**); in stage2, the number was 24 (22 preterm, 2 term) (**Figure 5f**); in stage3 infants, the rate of carriage was much lower, with only three infants (all preterm) potentially harbouring target species (**Figure 5g**). The change in prevalence of *K. oxytoca*-related species across the three stages based on the PanPhlAn analysis was consistent with the *K. oxytoca* abundance data generated with Centrifuge (**Figure 5b**).

The pangenome accessory genes were used in PCA to define which species strains detected by PanPhlAn belonged to. In stage1, samples from six preterm infants (P10111, P10141, P10301, P10451, P11292, P12121) and one term infant (P30221) harboured *K. michiganensis* (**Figure 5e**). In stage2, samples from four preterm infants (P10471, P10472, P11311, P11701) harboured *K. oxytoca*, while those from 14 preterm infants (P10071, P10231, P10301, P10441, P10451, P10501, P10601, P11151, P11202, P11291, P11292, P12121, P12641, P12651) and one term infant (P30221) harboured *K. michiganensis* (**Figure 5f**). Four samples (P11351, P12621, P20241, P30141) could not be assigned a species, while the sample from P11981 located close to *K. oxytoca* (**Figure 5f**). Similarly, two of the stage3 samples from P12221 and P12651 carried *K. michiganensis*, while P11981 located near *K. oxytoca* (**Figure 5g**).

### Recovery of *K. oxytoca* and *K. michiganensis* MAGs from metagenomes

Since the abundance of *K. oxytoca*-related species was considerable in some infant samples, we attempted to obtain high-quality MAGs directly from these metagenomes and to assign them to the species. The metagenomic samples were checked and their reads aligned against those of the 167- genome database; reads that mapped were extracted and assembled as ‘original MAGs’. The genome sizes of the 167 genomes ranged from 5.72 Mb to 7.23 Mb (mean 6.35 Mb), thus a genome size of at least 5.5 Mb was used to define a likely complete MAG. After assembly of the reads that mapped to our database, MAGs were generated from the stage 1 (_s1), stage2 (_s2) and stage3 (_s3) samples. ANI and phylogenetic analyses showed these MAGs to be *K. michiganensis* (P10301_s1, P10451_s1, P11292_s1, P12121_s1, P30221_s1, P10071_s2, P10301_s2, P10441_s2, P10451_s2, P10501_s2, P10601_s2, P11151_s2, P11202_s2, P11291_s2, P11292_s2, P12121_s2, P12641_s2, P12651_s2, P30221_s2, P12221_s3 and P12651_s3; mean ANI with GCA_002925905 of 98.61 ± 0.66 %) or *K. oxytoca* (P10472_s2, P11311_s2, P11701_s2, P11981_s2; mean ANI with GCA_900083895 of 99.30 ± 0.26 %) [34]. No *K. grimontii* MAGs were recovered from any samples.

Prior to checking the completeness and contamination of the *K. michiganensis* and *K. oxytoca* MAGs, contigs <500 nt in length were removed from the assemblies. A high-quality MAG requires a >90 % genome completeness with contamination <5 % [42]. According to CheckM results, three stage1 MAGs (P10301_s1, P10451_s1, P30221_s1), seven stage2 MAGs (P10441_s2, P10451_s2, P10501_s2, P10601_s2, P11291_s2, P11292_s2, P30221_s2) and one stage3 MAG (P12221_s3) were of high quality. The rest of the MAGs were ≥90 % complete, but were contaminated (e.g. P12621_2 contained 274.59 % contamination). Thus, we attempted to decontaminate the MAGs using a Diamond BLAST-based approach.

Since we already knew the species each MAG belonged to from PCA and PanPhlAn analyses, scaffolds were mapped against the relevant species-specific genome database under different minimum BLAST identity to report alignments, which were used to generate ‘cleaner’ MAGs. **Supplementary Figure 4** shows the change in genome completeness and the percentage contamination of stage2 MAGs when different blast identities were applied. The changes were negligible for those MAGs with high-quality-level completeness and lacking contaminants even when the cut-off was set at 99 %. For contaminated MAGs, the percentage contamination decreased markedly as the BLAST identity became stricter and reduced to the bottom when all scaffolds in that MAG could be aligned with 100 % identity. However, 100 % was not an appropriate threshold as the genome completeness was affected greatly at this point (**Supplementary Figure 4**). Instead, a cut-off of 99 % was used to decontaminate MAGs because 13 high-quality level MAGs and 5 medium-quality level MAGs could be obtained when using this identity threshold. Stage2 MAGs that passed PanPhlAn, PCA and ANI analysis reached at least reach medium-quality level using a 99 % identity threshold. This cut-off was also suitable for stage1 MAGs, the quality of which was high. However, for stage3 MAGs, the percentage contamination from P11981_s3 decreased to medium-quality level only at 100 % identity, at which time the genome completeness fell down to 42.40 %. After evaluating their genome completeness and contamination levels, a total of 25 MAGs (**Table 2**) were assessed further.

The presence of tRNAs for the standard 20 amino acids and rRNA was examined as a secondary measure of genome quality. A high-quality MAG requires at least 18 of the 20 possible amino acids [42]. P11981_s2 (16 aa) and P12651_s3 (17 aa) had to be classified as medium-quality MAGs even though their genome completeness and contamination reached high-quality levels. 16S rRNA genes were detected in all MAGs except P11151_s2, which was subsequently classified as medium quality. Taking mandatory genome information into consideration [42], a total of 19 high-quality and six medium-quality MAGs were recovered (**Table 2**); the sequences of these MAGs are provided in a **Supplementary File** (MAGs.zip). All of the MAGs had ≥15 standard tRNAs. High-quality MAGs had tRNAs that encoded an average of 19.6 (0.7) of the 20 amino acids, some of them even had a tRNA that encodes an additional amino acid SeC, while medium-quality MAGs had 18 (1.4) basic amino acids encoded by tRNAs. High-quality MAGs consisted of ≤500 scaffolds in 52.6 % of cases (mean 600) and had an average N50 of 121 kb, while only one medium-quality MAG comprised ≤500 scaffolds (mean 1174) and the average N50 was less than half of that of high-quality MAGs (48.3 kb).

### Genotyping of MAGs

Comparison of the sequences of the MAGs showed each infant harboured a different strain of *K. oxytoca* (**Figure 6a**) or *K. michiganensis* (**Figure 6b**). In infants where MAGs were recovered across different life stages, the MAGs were highly similar to one another (**Figure 6b**). Similar to what we had seen with our isolates, the MAGs encoded a range of β-lactamase and virulence genes (**Supplementary Figure 5**). It was also notable that two of the MAGs (*K. michiganensis* 10071_s2, *K. oxytoca* 10472_s2) encoded *mcr-9* (perfect match), a plasmid-mediated colistin resistance gene and phosphoethanolamine transferase. However, as noted above for our isolate work, presence of the aforementioned genes in MAGs does not mean they were functionally active in the infants’ GI tracts. All the *K. michiganensis* MAGs encoded the siderophore enterobacterin, along with all but one (11981_s2) of the *K. oxytoca* MAGs. The allantoinase gene cluster associated with liver infection was detected in the four *K. oxytoca* MAGs, but only a third of the *K. michiganensis* MAGs. We only detected this cluster in the *K. grimontii* isolates we recovered (**Figure 3b**).

**Figure 6.**
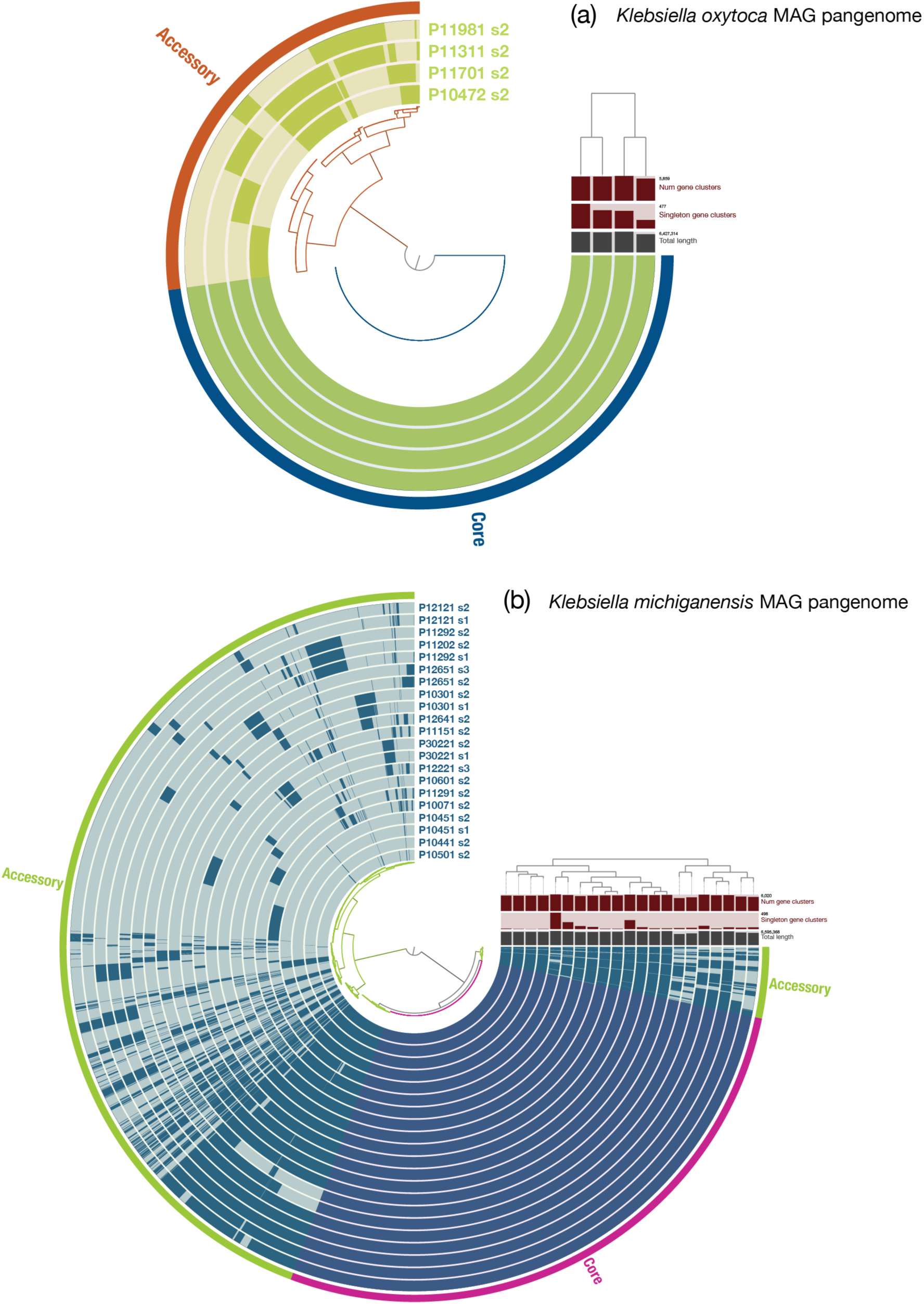
Anvi’o representation of the MAGs recovered from the metagenomes of infants included in the study of Ward *et al.* [32]. (a) *K. oxytoca*. (b) *K. michiganensis*. It is notable that MAGs recovered from different life stages from the same infant (e.g. 10301_s1, 10301_s2) are highly similar to one another.

MLST analysis assigned 10 MAGs to eight known STs (**Table 2**). We believe insufficient sequence coverage meant we were unable to ST more MAGs: e.g. *gapA* of MAG P11981_s2 aligned exactly with *gapA* allele 2 sequence but it was only partial, leaving an incomplete match.

The STs of all genomes in the curated genome dataset were also identified (**Supplementary Table 4**). Due to our limited understanding of *K. oxytoca*-related species, many combinations of alleles have not been assigned corresponding STs yet, especially for the newly described species *K. grimontii* [61]. MLST identification showed that 54/64 *K. oxytoca sensu stricto* strains had known STs, with some STs being more dominant than others. ST2, the most prevalent ST and represented by 15 strains, belonged to CC2. ST18 and ST19 were also in CC2, with both represented by two strains. ST199 and ST176 were the second and third most common STs, respectively. Among *K. michiganensis* strains, 48/76 of them could be assigned a known ST and comprised 21 distinct STs, with nine represented by more than one isolate. ST11, ST27, ST50, ST85, ST143 and ST202 were the most frequent STs, all of which were represented by at least four strains. *K. michiganensis* #108 was ST157. Only 5/27 *K. grimontii* strains could be assigned an ST (#83 – ST72; #99 – ST76; 10-5250 – ST47, GCA_000247915.1; 1148_KOXY – ST186, GCA_001052235.1; M5al – ST104, GCA_001633115.1).

## SUMMARY/CONCLUSIONS

*Klebsiella* spp. encode numerous virulence and antibiotic resistance genes that may contribute to the pathogenesis of NEC and LOS. In this study, we characterized nine *K. pneumoniae*, three *K. grimontii* and two *K. michiganensis* strains isolated prospectively from the faeces of a UK cohort of preterm infants, and have shown these gut isolates are able to reside and persist in macrophages, suggesting they can evade the immune system. These isolates will be used in future studies aiming to replicate aspects of NEC and sepsis in model systems to confirm the role of *Klebsiella* spp. in these diseases.

We have shown that mis-annotated genomes are being used in bioinformatics tools routinely used to characterize the human gut microbiome. By using a carefully curated dataset to undertake metapangenome analyses of the closely related species *K. oxytoca*, *K. michiganensis* and *K. grimontii*, we have demonstrated that *K. michiganensis* is likely to be more clinically relevant to a subset of preterm infants than *K. oxytoca*. Identity of publicly available genomes should be confirmed upon download and linked to accurate taxonomic frameworks prior to analyses of data, especially when attempting to identify and type closely related species in metagenomic data.

## Supporting information

MAG sequences

## ACKNOWLEDGEMENTS

Infrastructure support was provided by the National Institute for Health Research (NIHR) Imperial Biomedical Research Centre (BRC). This work used the computing resources of the UK MEDical BIOinformatics partnership – aggregation, integration, visualisation and analysis of large, complex data (UK Med-Bio) – which was supported by the Medical Research Council (grant number MR/L01632X/1), CLIMB [76] and the Nottingham Trent University Hamilton High Performance Computing Cluster. TCB was funded by a University of Westminster Faculty PhD studentship, and a Research Visit Grant from the Microbiology Society (RVG16/3). LH is a member of the ESGHAMI study group (https://www.escmid.org/research_projects/study_groups/host_and_microbiota_interaction/). This work was funded via a Wellcome Trust Investigator Award to LJH (100/974/C/13/Z), and support of the BBSRC Norwich Research Park Bioscience Doctoral Training Grant (BB/M011216/1, supervisor LJH, student CAG), and the Institute Strategic Programme Gut Microbes and Health BB/R012490/1, and its constituent project(s) BBS/E/F/000PR10353 and BBS/E/F/000PR10356 to LJH. We thank neonatal research nurses Karen Few, Hayley Aylmer and Kate Lloyd for obtaining parental consents and for collecting the samples at NNUH with the kind assistance of the clinical nursing team. We thank Matthias Scholz for his help in explaining the theory behind PanPhlAn to YC and advice on how to set appropriate parameters.

LH and LJH conceived and designed the study. All authors contributed to the writing of the manuscript. TCB did the isolation and initial characterization work, and isolated DNA from bacteria. YC did bioinformatics associated with genome analyses. OL did the anvi’o work. TCB, YC and OL were supervised by LH. LJH is chief investigator for the preterm clinical study from which samples were used, and PC is the clinical lead for the study. CAG performed 16S rRNA gene library preparation from faecal samples, and determined MIC values for strains. IO did macrophage assays. CZS did iron assays, and additional siderophore bioinformatics analysis. SC performed the 16S rRNA gene-associated bioinformatics, SP did 16S rRNA gene sequence analyses, and clinical database management. LJH supervised CA, IO, CZS, SC and SP.

---

**Supplementary Figure 1.** Genomic characterization of nine *K. pneumoniae* isolates recovered from neonate P008. (a) Change in number of new genes detected as the number of genomes increased. The number of new genes detected fell to almost 0 when a second genomes joined the first, and stayed at around 0 as further genomes were added to the analyses. (b, c) Change in total number of genes and conserved genes in the pangenome. As the number of genomes increased, the number of total genes (b) and conserved genes (c) is almost stable, with only negligible changes observed. (d) Number of BLASTP hits at different percentage identity. Identities of all hits are over 95 %, with the majority of them 100 % identical, providing more evidence to show the isolates from neonate P008 share the same genetic content. (e) Anvi’o representation of genomes of the isolates recovered from neonate P008. Differences between the isolates are due to the presence of single gene clusters in genomes, considered to be due to differences in genome coverage across the isolates.

**Supplementary Figure 2.** Anvi’o representation of genomes of the *K. pneumoniae* isolates recovered from different infants. The genomes are clearly different from one another, indicating each isolate represents a different strain of *K. pneumoniae*.

**Supplementary Figure 3.** Confirmation that the *K. oxytoca*-related isolates recovered from infants were strains of *K. michiganensis* and *K. grimontii*. (a) Heatmap generated with the R package heatmap.2() from FastANI [21] outputs. (b) Phylogenetic tree showing the placement of the genomes of the isolates among confirmed strains of *K. oxytoca*, *K. michiganensis* and *K. grimontii* [34]. FastTree v2.1.10 [37] was used to generate the tree from the core gene alignment produced by v3.12.0 (default settings) [36], with the tree visualized using FigTree v1.4.4 (http://tree.bio.ed.ac.uk/software/figtree/). In (a, b), affiliation of strains #88 and #108 with *K. michiganensis* is confirmed, while strains #80, #83 and #89 belong to *K. grimontii*.

**Supplementary Figure 4.** Genome quality assessment of MAGs. The change of genome completeness and percentage contaminations when different Diamond BLAST identities were applied to MAGs recovered from stage2 samples. Solid line represents completeness and dotted line represents contamination. An identity of 99 % is generally the most suitable cut-off to do decontamination as genome completeness is kept at a reasonable level but the removal of contaminants is effective.

**Supplementary Figure 5.** Summary of (a) antibiotic resistance and (b) virulence factor genes found in the MAGs recovered from the data of Ward *et al.* [32] by comparison of protein sequences with those of the CARD and VFDB, respectively. (a) Strict CARD match, not identical but the bit-score of the matched sequence is greater than the curated BLASTP bit-score cut-off; perfect CARD match, 100 % identical to the reference sequence along its entire length. Loose matches are not shown to avoid presenting false positives based on sequences with low homology and bit-scores below CARD BLASTP cut-off recommendations. (b) Identity (%), BLASTP reported only for those proteins sharing >70 % identity and 90 % query coverage with VFDB protein sequences.

