## Supplementary material for "Preterm infants harbour diverse *Klebsiella* populations, including atypical species that encode and produce an array of antimicrobial resistance- and virulence-associated factors": Figures

### Slide 1
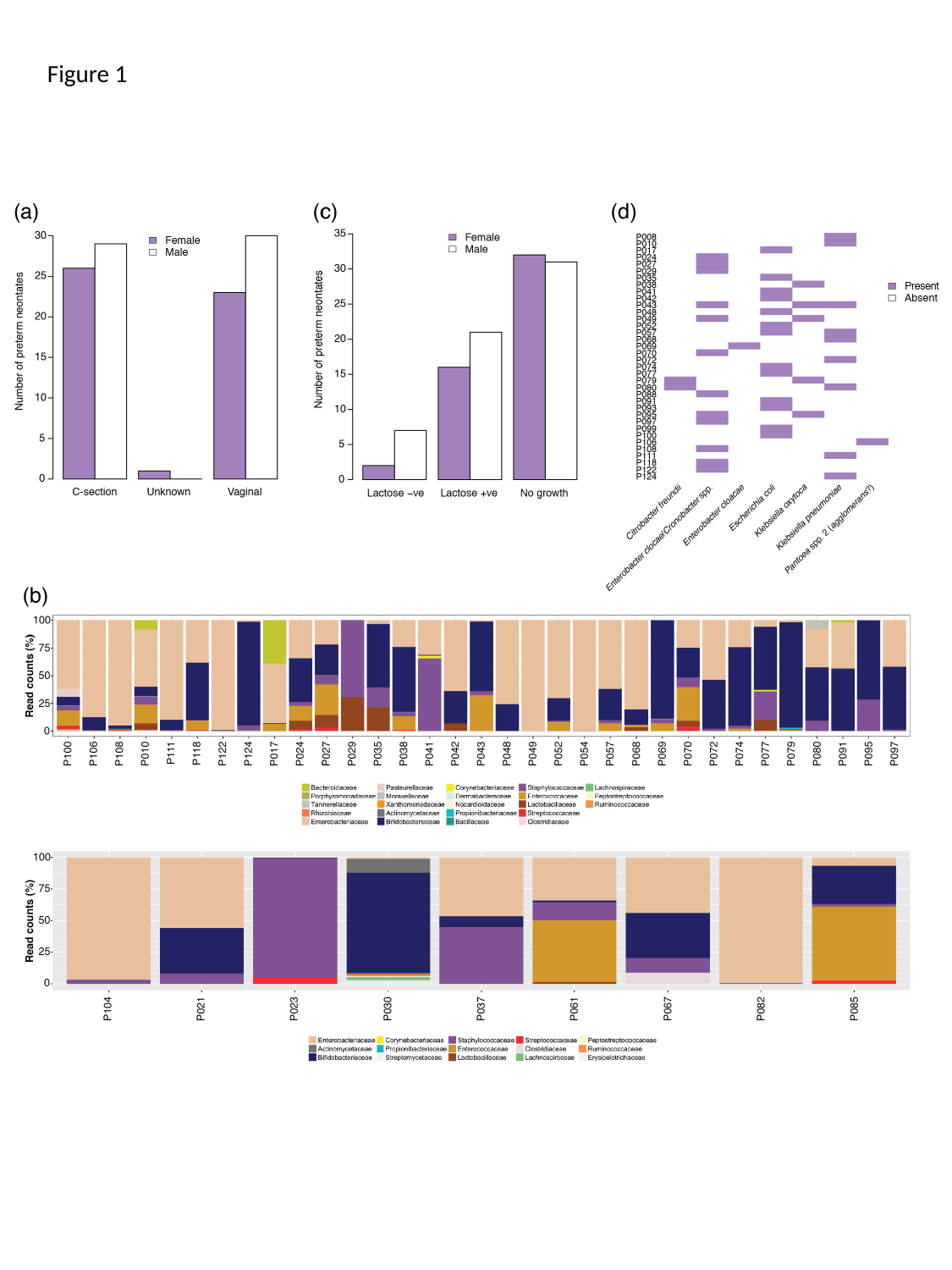

Figure 1

### Slide 2
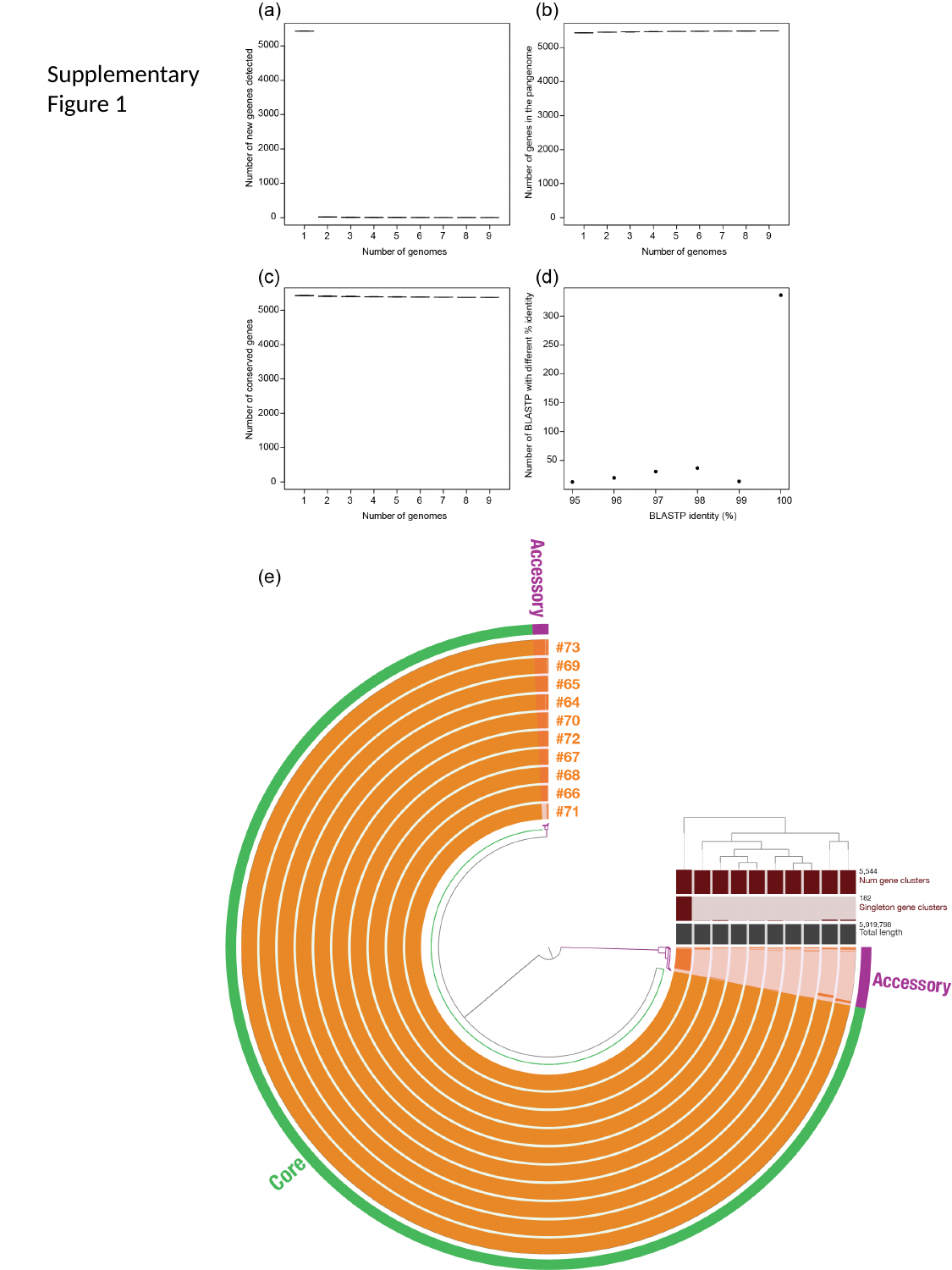

Supplementary Figure 1

### Slide 3
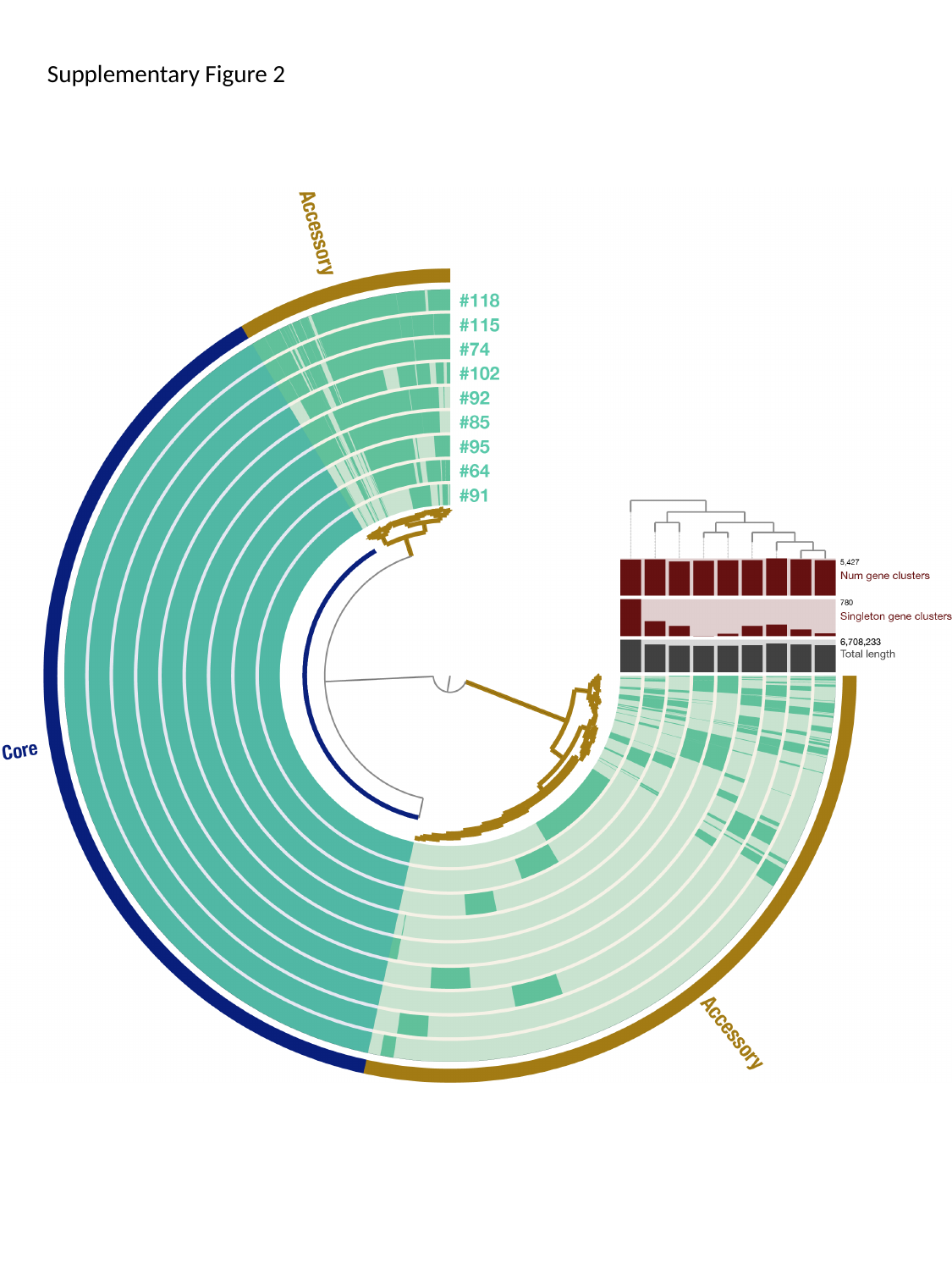

Supplementary Figure 2

### Slide 4
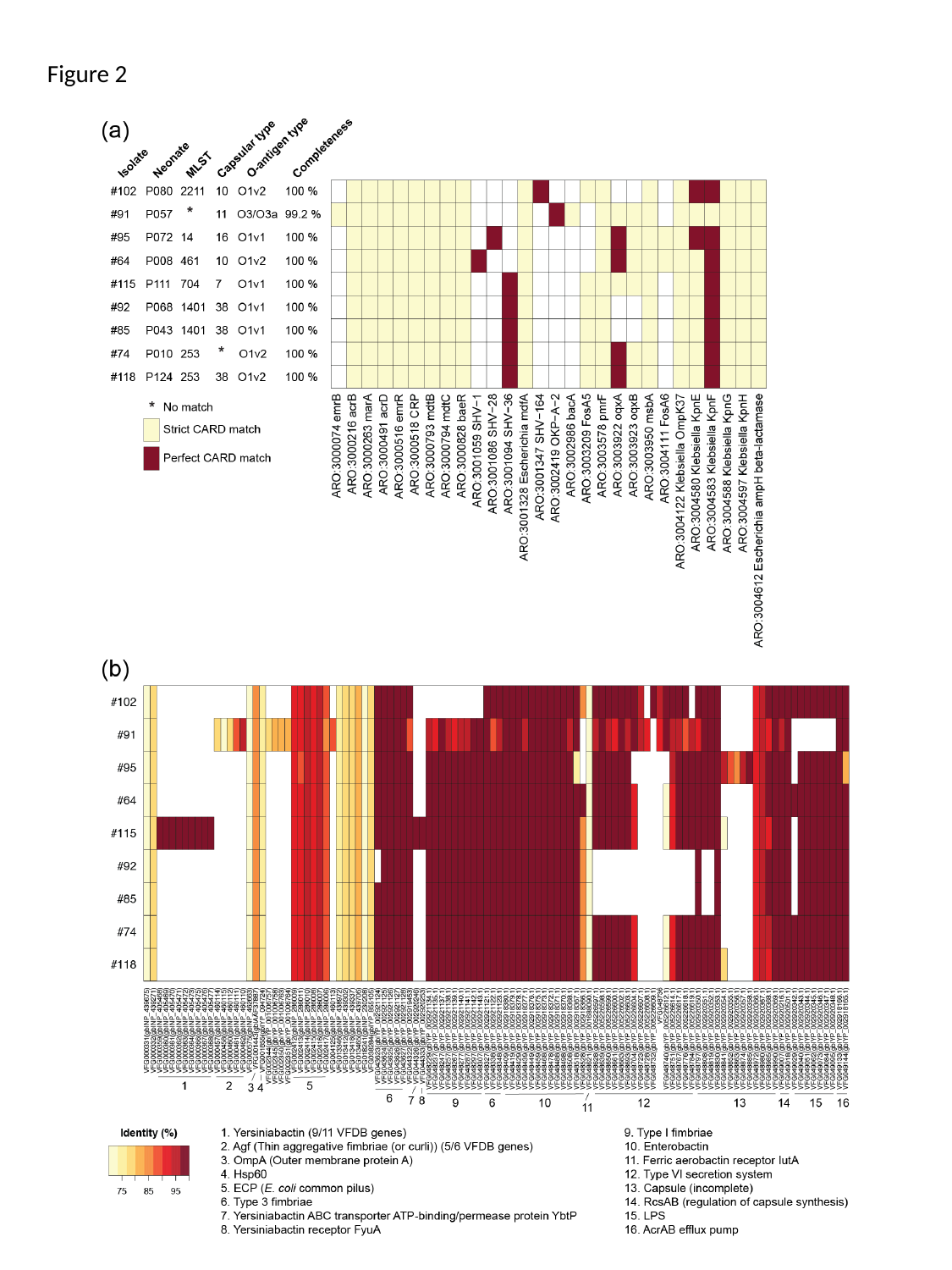

Figure 2

### Slide 5
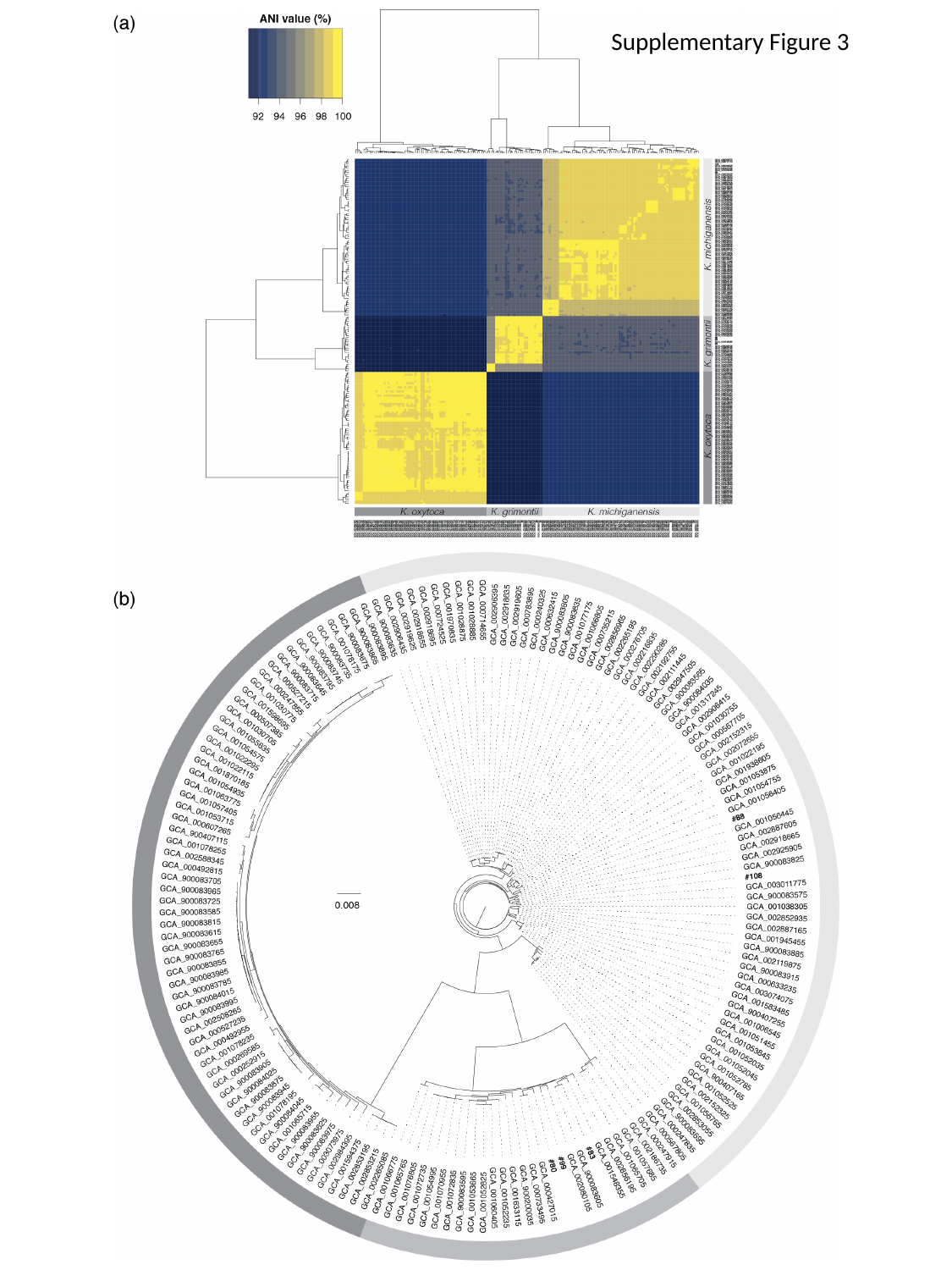

Supplementary Figure 3

### Slide 6
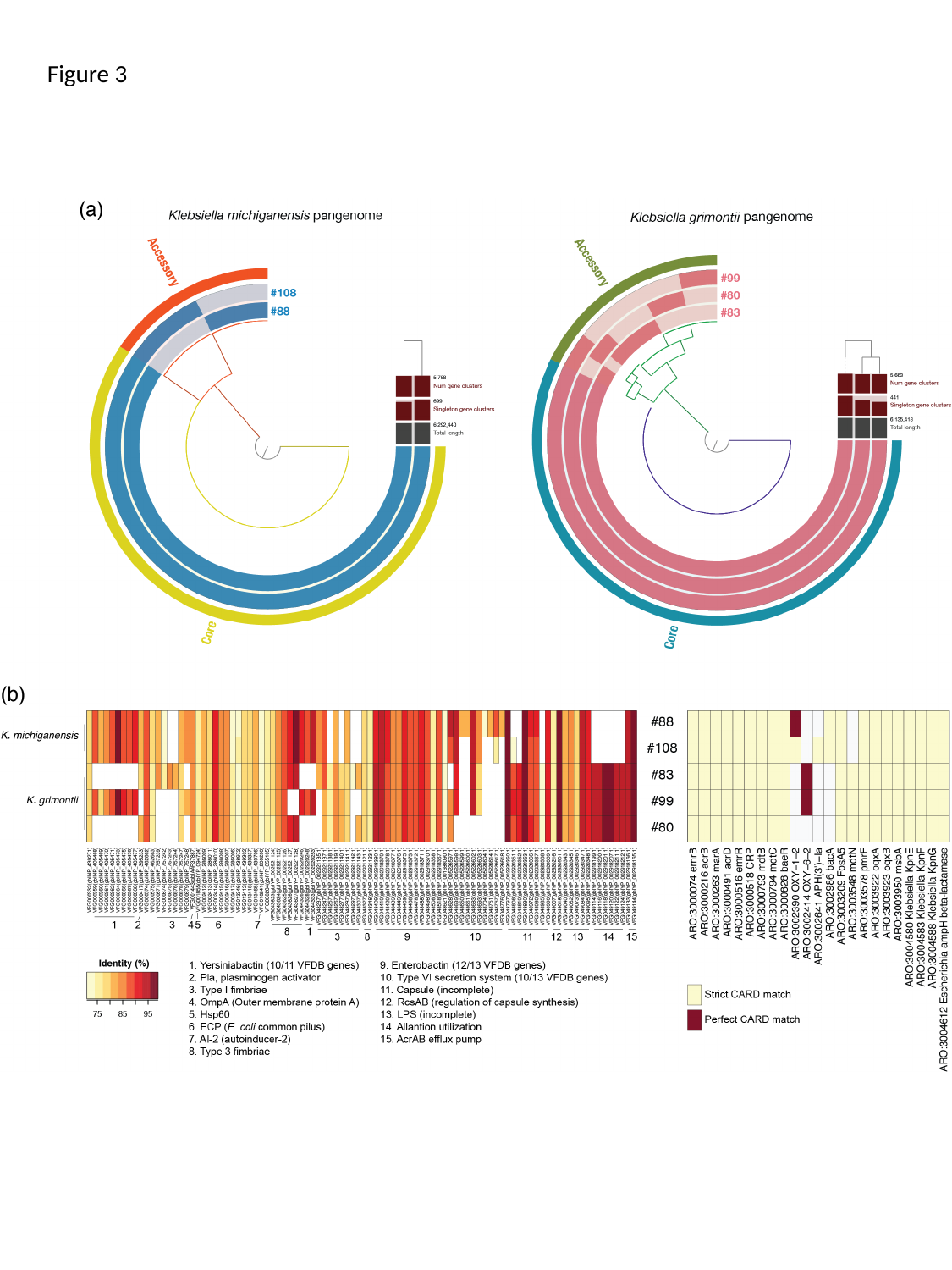

Figure 3

### Slide 7
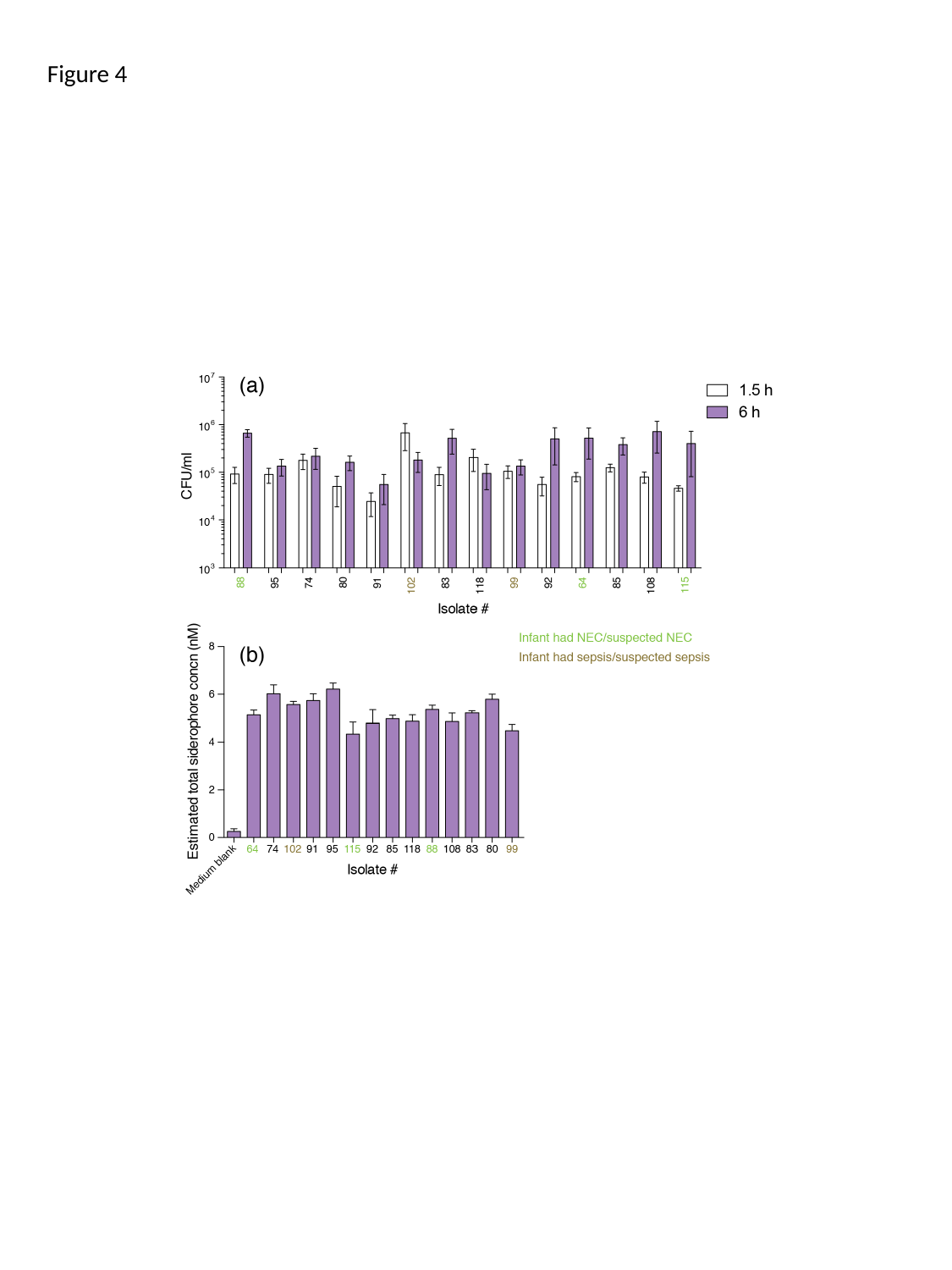

Figure 4

### Slide 8
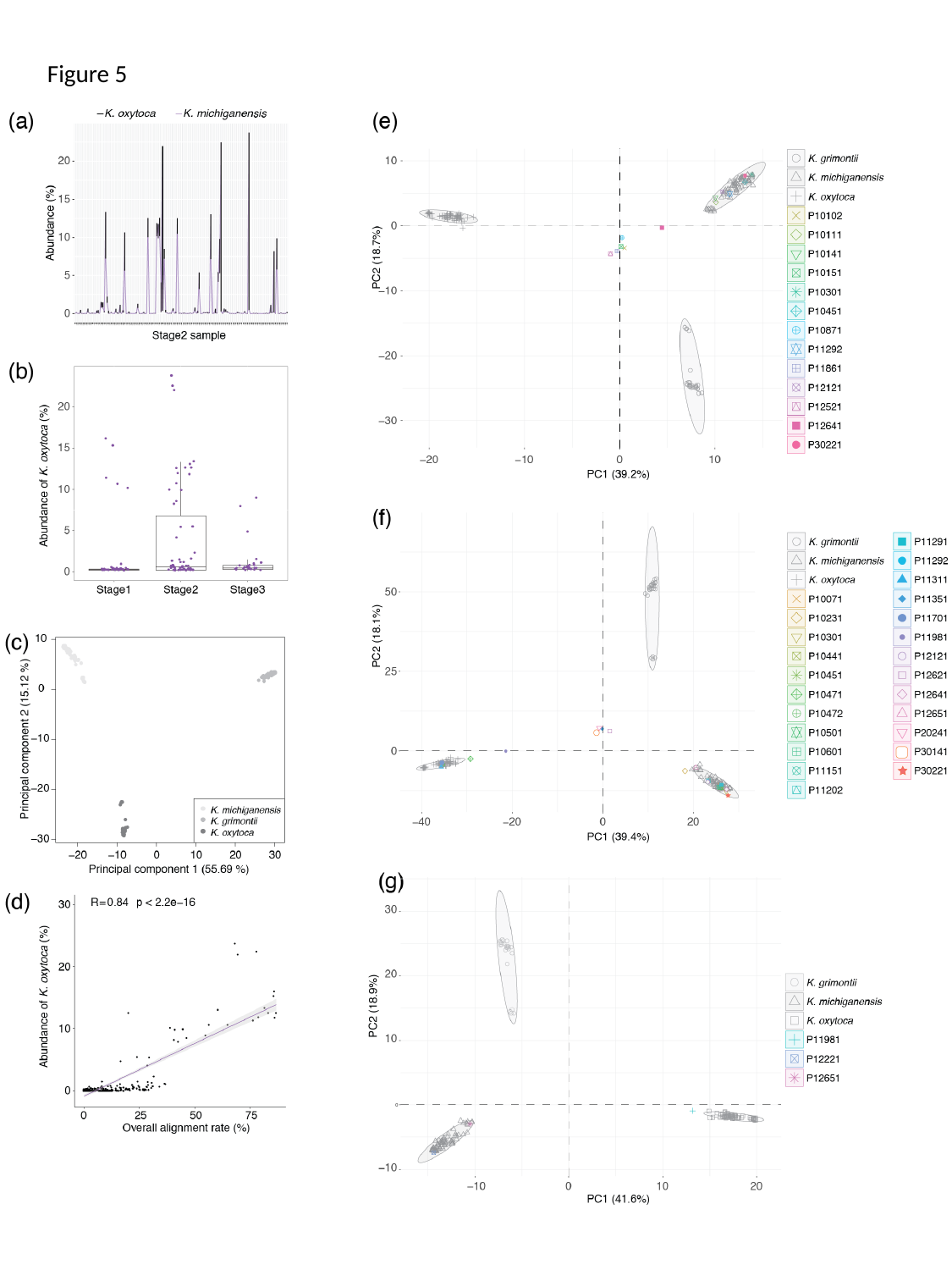

Figure 5

### Slide 9
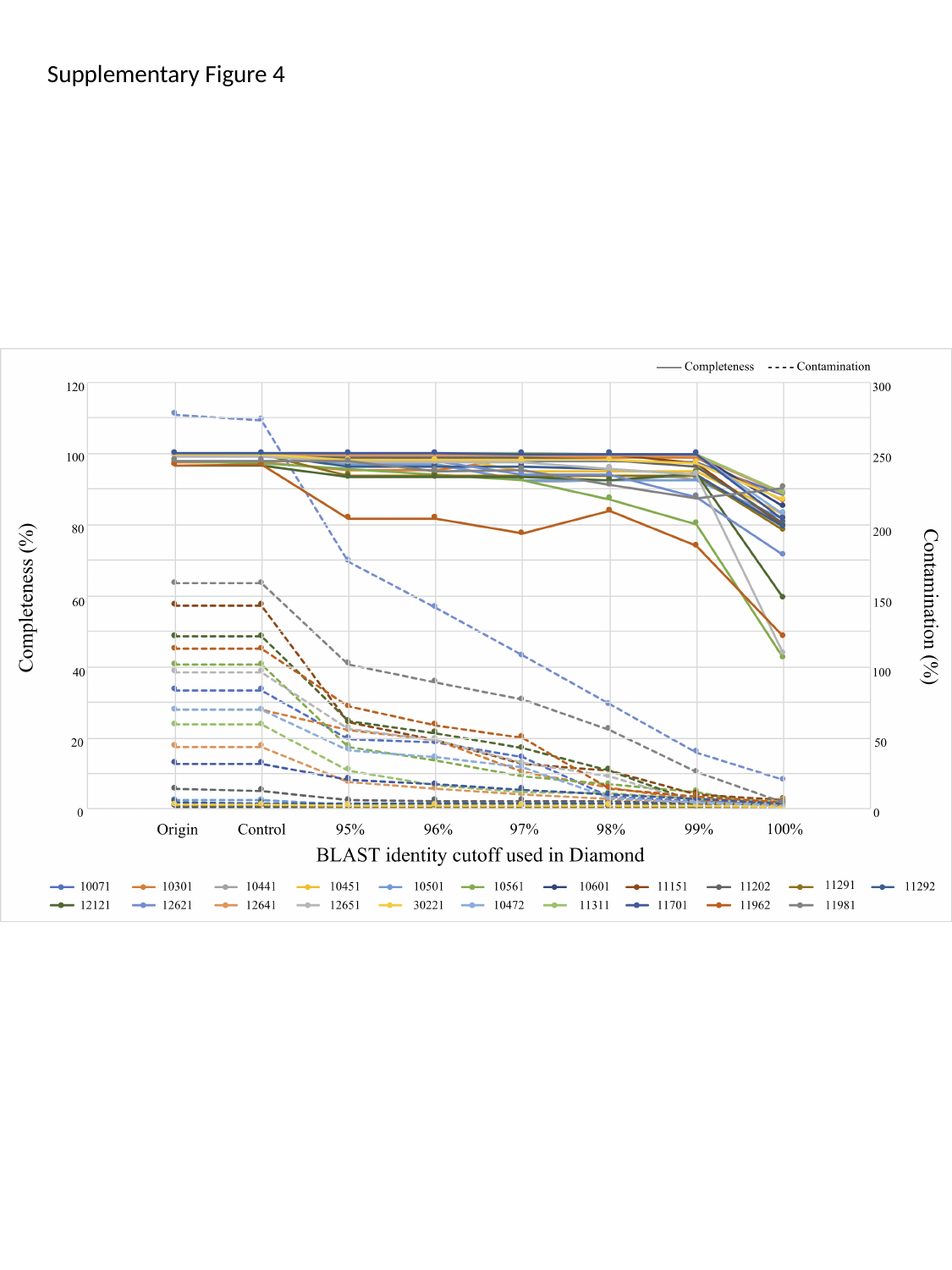

Supplementary Figure 4

### Slide 10
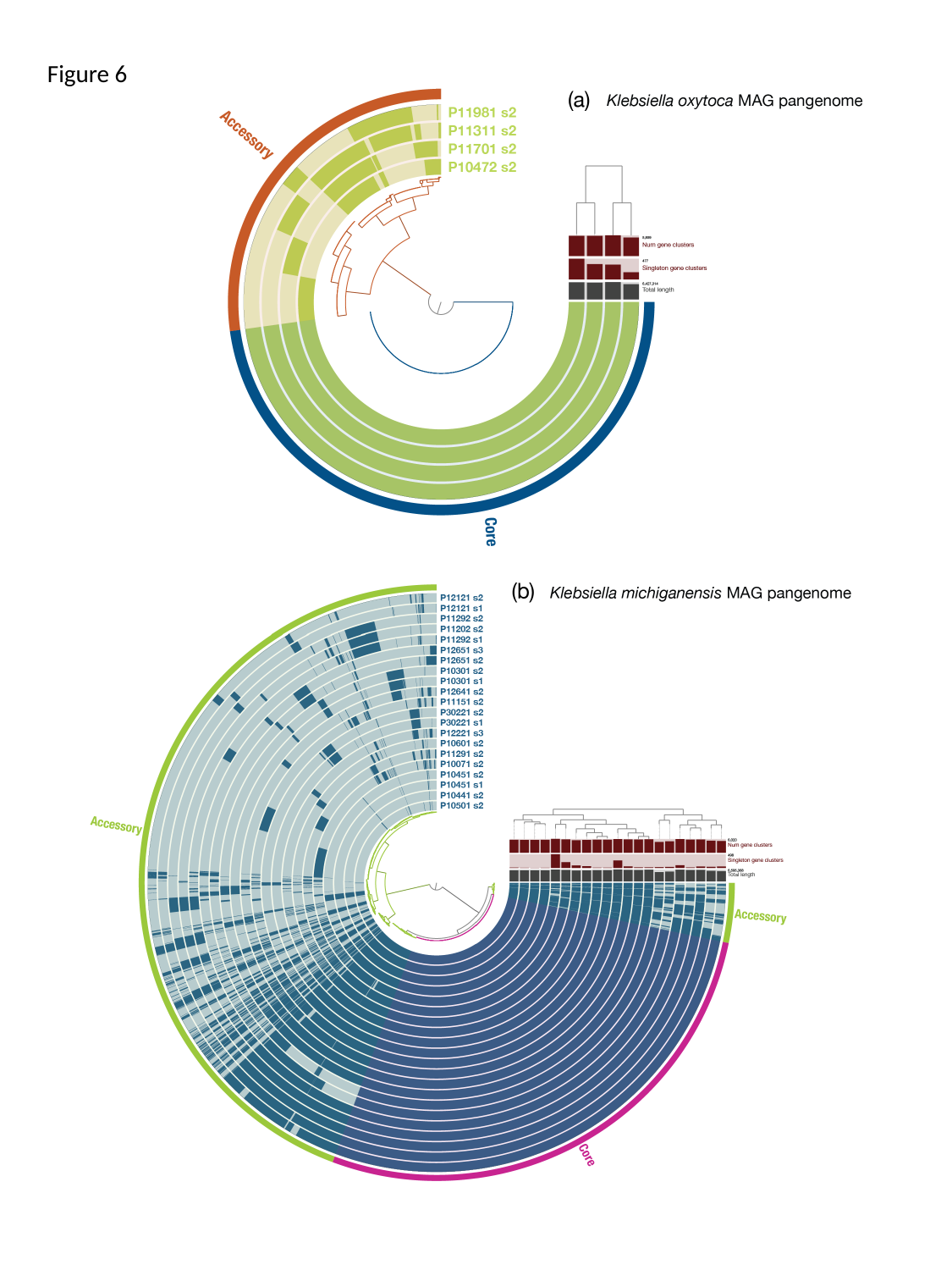

Figure 6

### Slide 11
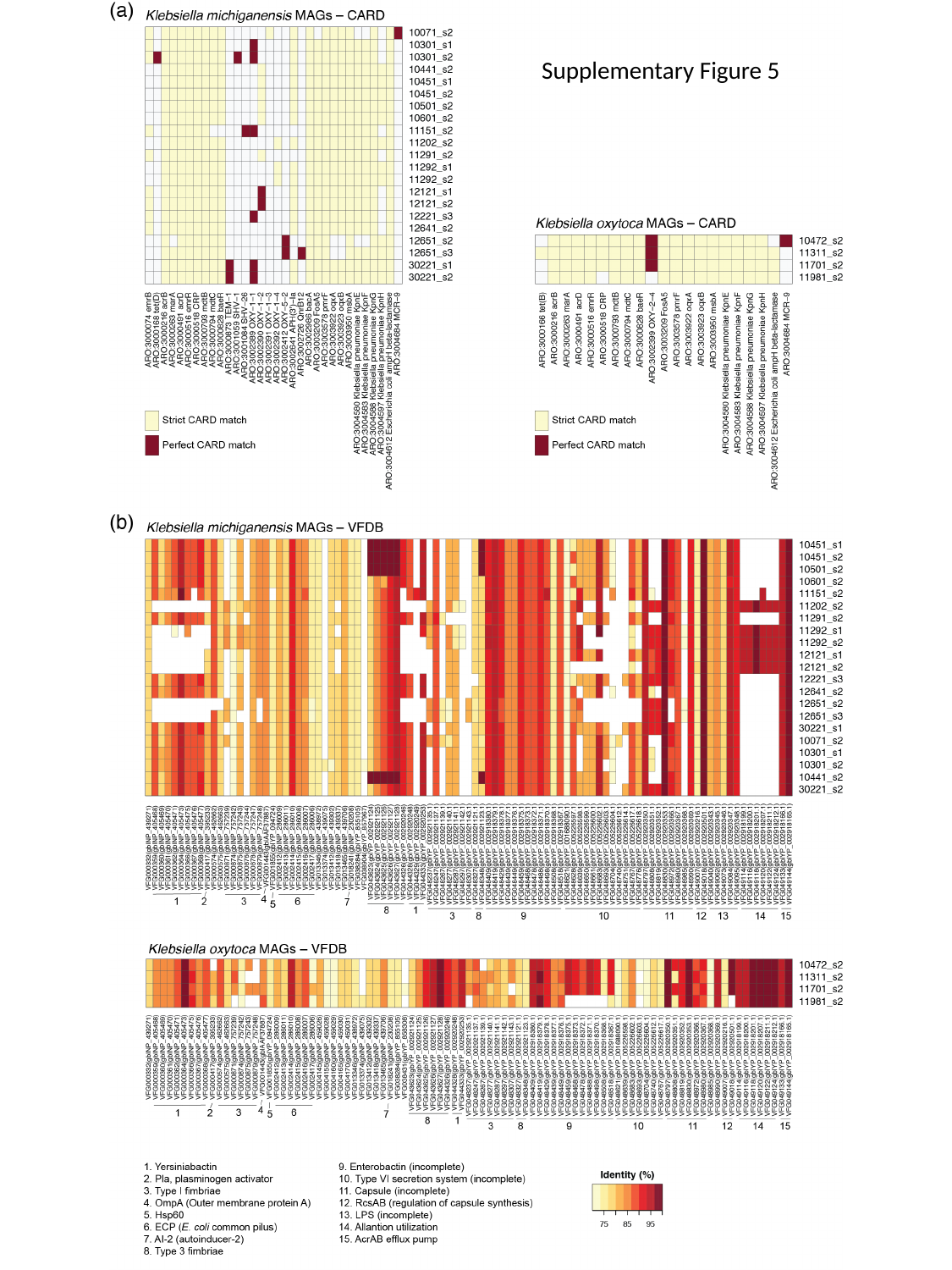

Supplementary Figure 5
